# The structure of the full catalytic cycle of *Vibrio cholerae* NFeoB

**DOI:** 10.1101/2025.10.06.680749

**Authors:** Kate Magante, Candice M. Armstrong, Mark Lee, Aaron T. Smith

## Abstract

The acquisition of iron is critical for the survival and the virulence of numerous infectious pathogens, and most bacteria acquire ferrous iron (Fe^2+^) by utilizing the ferrous iron transport (Feo) system. FeoB is the main component of this system, and its function is regulated by the soluble cytosolic domain, termed NFeoB. We have recently begun to define the structure and the mechanism of the Feo system from the bacterium *Vibrio cholerae*, the causative agent of the disease cholera. However, major structural gaps in our understanding of the nucleotide-promiscuous *V. cholerae* NFeoB still exist. In this work, we have determined several new X-ray crystal structures that reveal distinct snapshots of the *Vc*NFeoB domain in uncommon and unprecedented states, ultimately illuminating the full catalytic cycle of this NTPase. This work reveals important functional features of *Vc*NFeoB that may be leveraged and ultimately targeted to prevent the infectivity and the spread of cholera.

**Graphical Abstract:** 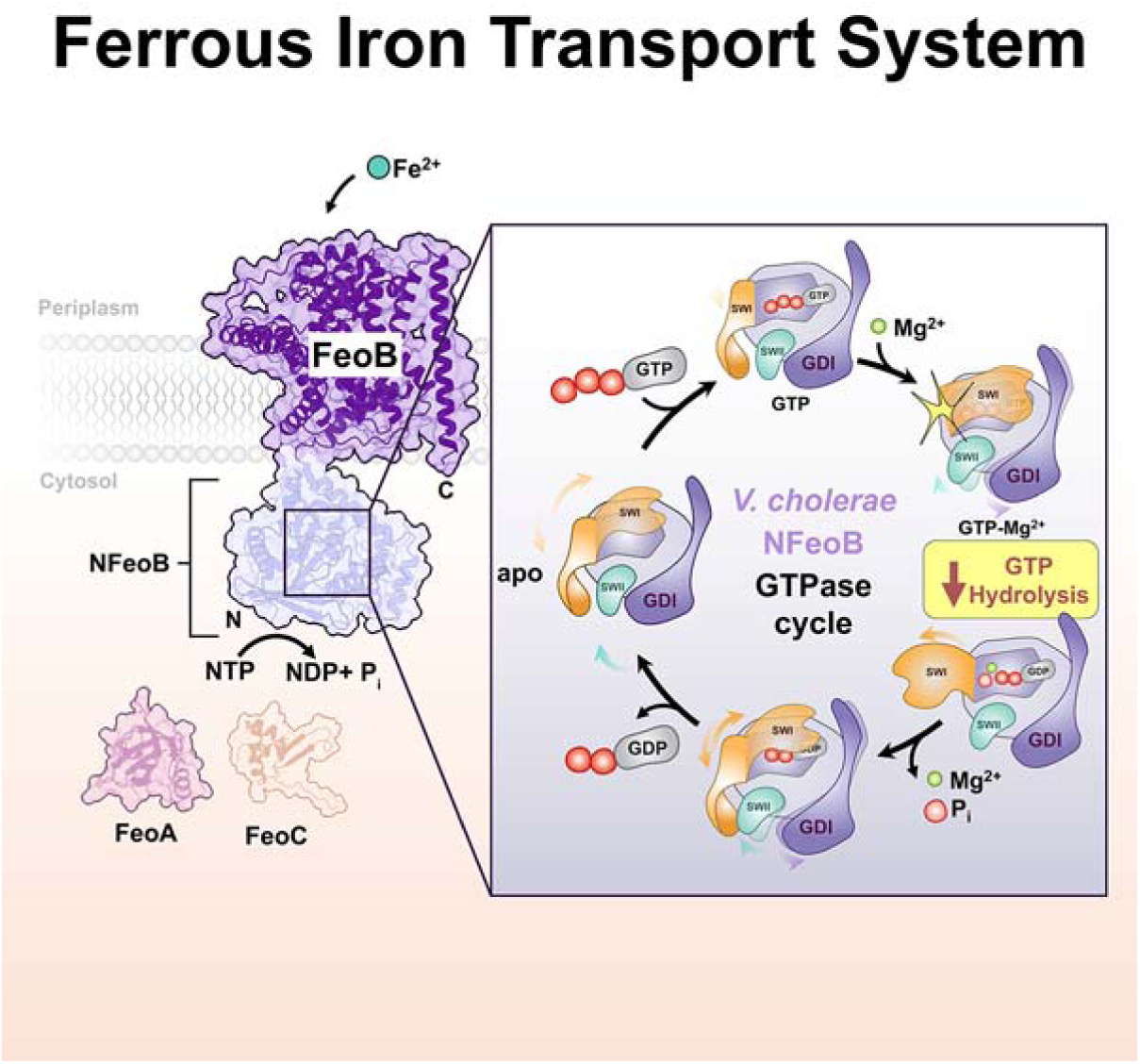

## INTRODUCTION

The acquisition of iron by organisms is critical, as this essential trace element functions as an obligate cofactor in numerous biological processes such as oxidative phosphorylation, DNA biosynthesis, cellular respiration, and even N_2_ fixation.^1–4^ Despite this importance, our understanding of how iron is obtained by microorganisms is especially understudied, leading to major gaps in our knowledge of how this element is linked to human diseases. For infectious bacteria within a host, distinct populations of both ferric (Fe^3+^) and ferrous (Fe^2+^) iron may exist under different physiological conditions and may be bound to different types of biomolecules.^5–7^ In O_2_-replete environments, the formation of insoluble iron hydroxides/oxides occurs spontaneously and represents a thermodynamic sink. To access soluble Fe^3+^, bacteria have adapted to scavenge ferric iron by using low molecular weight siderophore-based approaches.^8–10^ Pathogenic bacteria may also release proteins that will bind to either extracellular free Fe^3+^ ions/molecules, or will utilize transferrin/lactoferrin-binding proteins that will sequester Fe^3+^ from their respective polypeptides.^9, 11^ Infectious bacteria may also scavenge iron protoporphyrin IX (heme *b*) from erythrocytes and muscle tissue, if available.^7, 12–13^ While both siderophore-and heme-based Fe^3+^ acquisition strategies are actively studied by multiple research groups^14–18^ and have been the target of recent exploits to attenuate bacterial virulence^19–20^, our comparative understanding of bacterial Fe^2+^ acquisition remains in its infancy despite its important connection to human diseases.^11, 21–24^

Under acidic and/or anaerobic conditions, such as those found in the gut or within biofilms, iron is chiefly found in the reduced (Fe^2+^) form and is acquired by bacteria typically using the Feo system (Figure 1), a multi-protein complex that is also leveraged by numerous infectious bacteria to establish pathogenesis.^6, 21, 25–26^ While in most bacteria the *feo* operon encodes for only two proteins (FeoA and FeoB), the *feo* operon in clinically-relevant γ-proteobacteria (such as *Vibrio cholerae*, the pathogenic bacterium responsible for the diarrheal disease cholera) also encodes a third protein, FeoC.^6, 21, 25, 27–29^ FeoA and FeoC are small cytosolic proteins, while FeoB is a large transmembrane protein bound to an N-terminal soluble G-protein-like domain termed NFeoB.^6, 21, 25^ In most organisms, (N)FeoB binds and hydrolyzes GTP exclusively; however, in some organisms (such as *V. cholerae*) (N)FeoB is more promiscuous and may accommodate other nucleotides.^27, 30–33^ The status of nucleotide bound to the NFeoB region is hypothesized to control the opening and closing of the FeoB pore and therefore the translocation of Fe^2+^ into the bacterial cytosol.^21, 26^ Yet, how FeoA, FeoB, and/or FeoC function together to mobilize Fe^2+^ across a lipid bilayer remains unclear.

**Figure 1.**
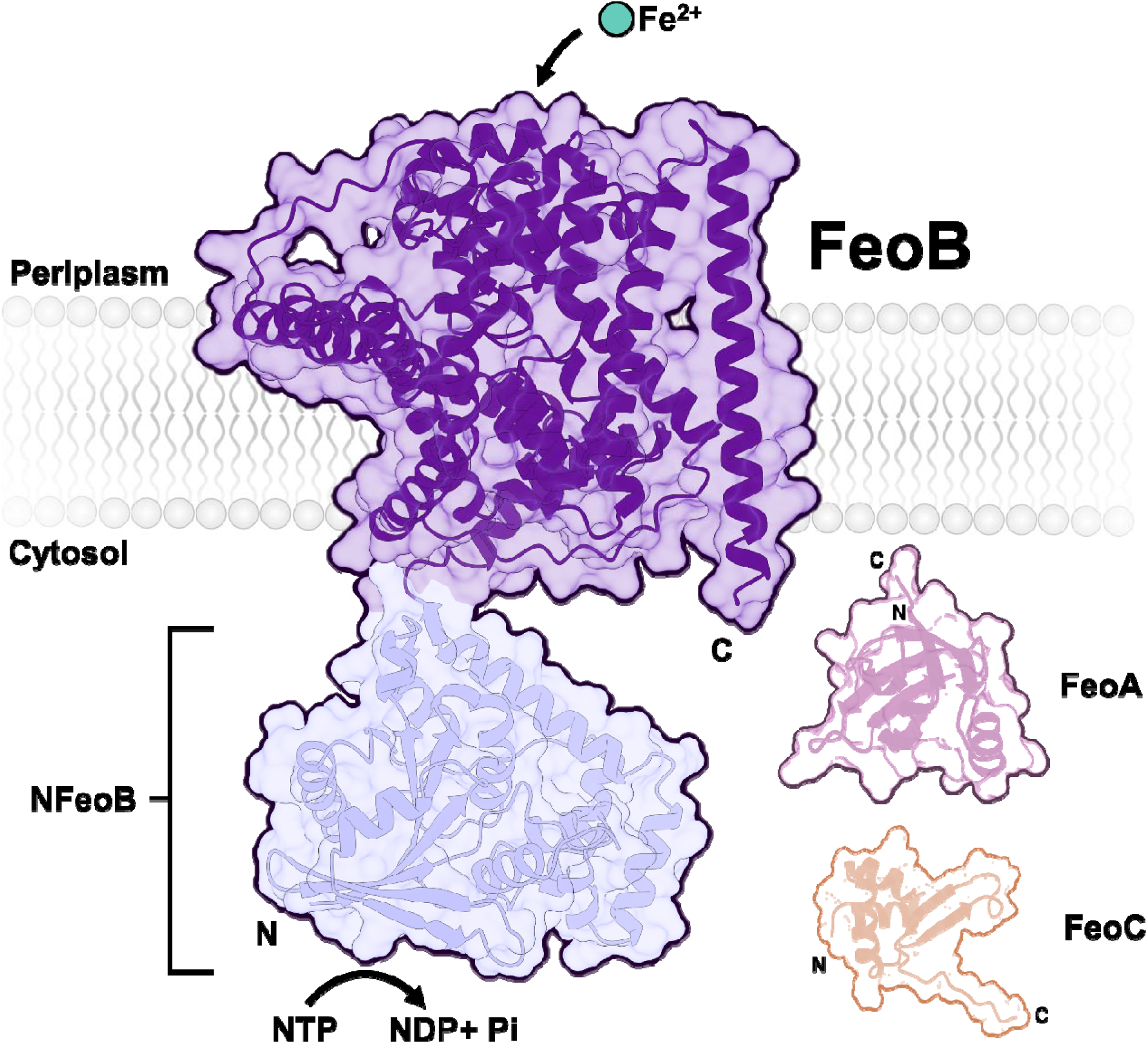
Cartoon representation of the tripartite Feo system in *Vibrio cholerae*. The *V. cholerae* Feo system is composed of three proteins: FeoA, FeoB, and FeoC. FeoA and FeoC are small cytosolic proteins, while FeoB is a transmembrane protein with a soluble N-terminal domain termed NFeoB. Structures of *Vc*NFeoB and *Vc*FeoC have been experimentally determined (PDB IDs 8VWL and 7U37, respectively), while the structures of *Vc*FeoA and intact *Vc*FeoB are AlphaFold models (AF-A0A655V7F8-F1-v4 and AF-A0A5Q6PH70-F1-v4, respectively). *V. cholerae* FeoB is one of several FeoBs known to be a promiscuous NTPase rather than a strict GTPase.

Extensive *in vivo* data using chiefly *V. cholerae* have shown that the Feo proteins form a complex that controls Fe^2+^ uptake^34–35^, and we have recently begun to define the structure of this complex. For example, we first determined the NMR structure of *Vc*FeoC in solution, revealing that FeoCs conserve their helix-turn-helix (HTH) motifs but not their [Fe-S] binding domains.^36–38^ Using recombinant approaches, we demonstrated that *Vc*FeoC interacts with full-length, intact *Vc*FeoB, and homology modeling suggested these protein-protein interactions are mediated by the FeoC HTH domain and occur within proximity of two key regulatory loops (known as Switch I and Switch II) in the *Vc*NFeoB cytosolic domain.^36^ We next determined to modest resolution the X-ray crystal structures of WT and variant *Vc*NFeoBs in both the apo and GDP-bound forms, revealing a canonical NFeoB G-protein fold despite the nucleotide promiscuity of *Vc*NFeoB.^31^ Using modeling and biophysical approaches, we proposed a framework for how NTP-agnostic FeoBs could function adaptively to ensure bacterial iron quotas across changing metabolic landscapes.^31^ However, we previously only determined the partial catalytic cycle of *Vc*NFeoB; in fact, the complete catalytic cycle (including a transition-state analog) of NFeoB has only ever been determined for a single thermophile (*Staphylococcus thermophilus*)^39–40^ but not for a clinically-relevant mesophile such as *V. cholerae*, representing a major gap in our understanding of the structure of this system.

In this work, we have determined the structural basis of the full catalytic cycle of *V. cholerae* NFeoB. Uniquely, all forms of *Vc*NFeoB crystallized in the *P_1_* space group with 4 molecules in the asymmetric unit (ASU), allowing us to visualize multiple poses of the protein under different conditions. Specifically, we have captured snapshots of *Vc*NFeoB in complex with the non-hydrolyzable GTP analog guanosine-5′-[(β,γ)-methyleno] triphosphate (GMP-PCP) with and without Mg^2+^, *Vc*NFeoB in complex with guanosine diphosphate (GDP) bound to AlF_3_ and Mg^2+^ (representing a transition-state analog), *Vc*NFeoB in complex with GDP and Mg^2+^ (post phosphate release), and *Vc*NFeoB in complex with GDP (post phosphate and Mg^2+^ release) at significantly improved resolution. Surprisingly, the structure of transition-state analog-bound *Vc*NFeoB reveals Switch I in an open conformation poised for phosphate release, in contrast with the transition-state analog-bound form of *St*NFeoB in which Switch I adopted a conformation covering the nucleotide-binding pocket. Moreover, our work shows that Mg^2+^ plays a critical role in mobilizing Switch I for nucleotide hydrolysis, which is further probed using isothermal titration calorimetry (ITC). When taken together, these results reveal the full GTP-hydrolysis cycle of *Vc*NFeoB and suggest important structural and functional features of *Vc*NFeoB that may be targeted to prevent the infectivity and spread of cholera.

## RESULTS

### Crystallization of the nucleotide-bound forms of VcNFeoB

To crystallize its entire catalytic cycle, we prepared highly pure *Vc*NFeoB similar to our prior approach.^31^ Briefly, we first expressed and purified an N-terminally (His)_6_-tagged SUMO fusion of the *Vc*NFeoB protein. After cleavage, tag separation, and polishing, the resultant *Vc*NFeoB was highly pure and chiefly monomeric in solution based on analytical size-exclusion chromatography and mass photometry (Fig. S1). This solution-state oligomerization is consistent with our prior analyses of recombinant forms of FeoA^41–42^, NFeoB^31^, FeoA fused to NFeoB^42^, FeoC^36–37^, and intact FeoB^32, 43^, which is also consistent with work on truncated and intact FeoB from other labs^44^. To generate the guanosine diphosphate (GDP)-bound and guanosine-5′-[(β,γ)-methyleno] triphosphate (GMP-PCP)-bound forms of *Vc*NFeoB in solution, we pre-incubated the protein with excess nucleotide (both with and without Mg^2+^) at room temperature for 2 hr prior to setting up crystal trays at 20 °C. To generate the GDP-Mg^2+^-AlF_3_-bound form of *Vc*NFeoB, we co-crystallized the GDP-bound form of *Vc*NFeoB with NaF and AlCl_3_ using an approach that previously generated the GDP-Mg^2+^-AlF_4_ ^-^-bound form of *Staphylococcus thermophilus* NFeoB (*St*NFeoB).^40^ Unlike the other nucleotide-bound forms of *Vc*NFeoB, and unlike *St*NFeoB, the structure of *Vc*NFeoB containing a transition-state analog was only observed in crystals formed at 4 °C and not at other temperatures, suggesting that the nucleotide hydrolysis transition state of a mesophilic NFeoB is highly dynamic at room temperature, at least in this crystal form. Nonetheless, the structures derived from these crystals (Table 1; Fig. S2) were combined with our previously-determined apo *Vc*NFeoB structure^31^ to elucidate the full GTP hydrolysis cycle catalyzed by *Vc*NFeoB.

**Table 1.** Data collection and refinement statistics for WT*Vc*NFeoB + GDP (PDB ID 9D8B), WT*Vc*NFeoB + GMP-PCP (PDB ID 9D8D), and WT*Vc*NFeoB + GDP +AlF_3_ (PDB ID 9PSD) all in the *P_1_* space group containing 4 molecules of *Vc*NFeoB in the asymmetric unit (ASU). Parentheses indicate the highest resolution shells.

| | WTVcNFeoB + GMP-PCP in $P_1$ space group (PDB ID 9D8D) | WTVcNFeoB + GDP + AlF<sub>3</sub> in $P_1$ space group (PDB ID 9PSD) | WTVcNFeoB + GDP in $P_1$ space group (PDB ID 9D8B) |
| --- | --- | --- | --- |
| **Data Collection** |  |  |  |
| Beamline | NSLS-II 17-ID-2 | NSLS-II 17-ID-2 | NSLS-II 17-ID-2 |
| Wavelength (Å) | 0.97936 | 0.97934 | 0.97936 |
| Space group | $P_1$ | $P_1$ | $P_1$ |
| Cell Dimensions |  |  |  |
| $a, b, c$ (Å) | 53.98, 59.67, 89.43 | 54.24, 59.08, 89.63 | 54.05, 59.02, 89.48 |
| $\alpha, \beta, \gamma$ (°) | 93.59, 92.61, 113.77 | 94.44, 92.37, 113.54 | 94.39, 92.98, 113.21 |
| Resolution (Å) | 88.99-2.11 | 34.52-2.00 | 32.65-1.82 |
| $R_{\text{merge}}$ | 0.051 (0.642) | 0.153 (1.059) | 0.040 (0.924) |
| $CC_{1/2}$ | 0.990 (0.557) | 0.986 (0.329) | 0.997 (0.452) |
| $I/\sigma(I)$ | 9.4 (0.6) | 4.0 (1.4) | 4.2 (0.5) |
| Completeness (%) | 97.6 (92.0) | 91.9 (97.1) | 93.9 (94.4) |
| **Refinement** |  |  |  |
| Resolution (Å) | 31.81-2.11 (2.15-2.11) | 32.72-2.00 (2.03-2.00) | 30.83-1.82 (1.84-1.82) |
| No. reflections | 56,827 (2669) | 62,614 (2969) | 84,916 (2770) |
| $R_{\text{work}}$ | 0.196 | 0.188 | 0.203 |
| $R_{\text{free}}$ | 0.246 | 0.242 | 0.247 |
| No. atoms/molecules |  |  |  |
| Protein | 7,890 | 8,021 | 7,826 |
| Mg<sup>2+</sup> | 2 | 2 | 0 |
| GDP | 0 | 4 | 4 |
| GMP-PCP | 4 | 0 | 0 |
| AlF<sub>3</sub> | 0 | 1 | 0 |
| Waters | 168 | 618 | 228 |
| Glycerol | 1 | 3 | 7 |
| Average $B$-factors (Å<sup>2</sup>) | 66.92 | 29.00 | 50.99 |
| R.m.s. deviations |  |  |  |
| Bond lengths (Å) | 0.008 | 0.007 | 0.009 |
| Bond angles (°) | 1.00 | 0.96 | 1.14 |
| Ramachandran plot |  |  |  |
| Favored | 98.4 % | 98.3 % | 98.9 % |
| Allowed | 1.6 % | 1.7 % | 1.1 % |
| Outlier | 0.0 % | 0.0 % | 0.0 % |

### The apo state of VcNFeoB reveals dynamic Switch I and Switch II regions in the absence of nucleotide

The apo form of *Vc*NFeoB is the starting point of the nucleotide hydrolytic cycle, and our nucleotide- and Mg^2+^-free structure of *Vc*NFeoB reveals key regions of the protein that are dynamic prior to nucleotide binding. The overall structure of the apo *Vc*NFeoB domain (PDB ID 9BA6; 2.38 Å resolution) can be seen in Fig. 2a and closely resembles the structure of other native and variant NFeoBs from Gram-negative γ-proteobacteria.^44–47^ Germane to the results in this work, several key regions are highlighted in the structure and the sequence of *Vc*NFeoB (Fig. 2a,b), such as the G1-G5 motifs (responsible for the recognition and binding of guanine nucleotides in most GTPases)^6, 48–49^, the Switch I and Switch II regions (critical for nucleotide hydrolysis and hypothesized to sense and to communicate nucleotide status in FeoB)^6, 48^, and the guanine-dissociation inhibitor domain (connected directly to the transmembrane region of intact FeoB in the primary amino acid sequence and known to prevent GDP-to-GTP exchange^6, 47, 50^). Interestingly, unlike other crystallized forms of apo NFeoB (*e.g.*, *Escherichia coli* NFeoB^45^, *Klebsiella pneumoniae* NFeoB^46^, *Legionella pneumophila* NFeoB^47^), apo *Vc*NFeoB displays dynamics in both the Switch I and Switch II regions in the absence of nucleotide (Fig. 2b; Fig. S3). In one form in the asymmetric unit (ASU; chain A), Switch I is nearly completely ordered, with the exception of a short turn from Trp32 to Gly34; however, this ordered Switch I correlates with a more disordered Switch II in chain A, as a more extensive loop comprising Gly65 to Asn69 could not be observed in the electron density (similar to the apo structures of *Kp*NFeoB and *Lp*NFeoB)^46–47^ (Table 2). In striking contrast, in chain B, Switch I residues stretching from Lys26 through Glu38 (*i.e.*, the majority of Switch I) are completely dynamic and disordered, while the entirety of Switch II is observed in the electron density (Fig. 2b; Table 2; Fig. S3). This interesting result contrasts with previous apo structures of NFeoB but logically suggests that these two regions are highly dynamic in the absence of nucleotide, likely communicating the nucleotide status to the rest of the protein.

**Figure 2.**
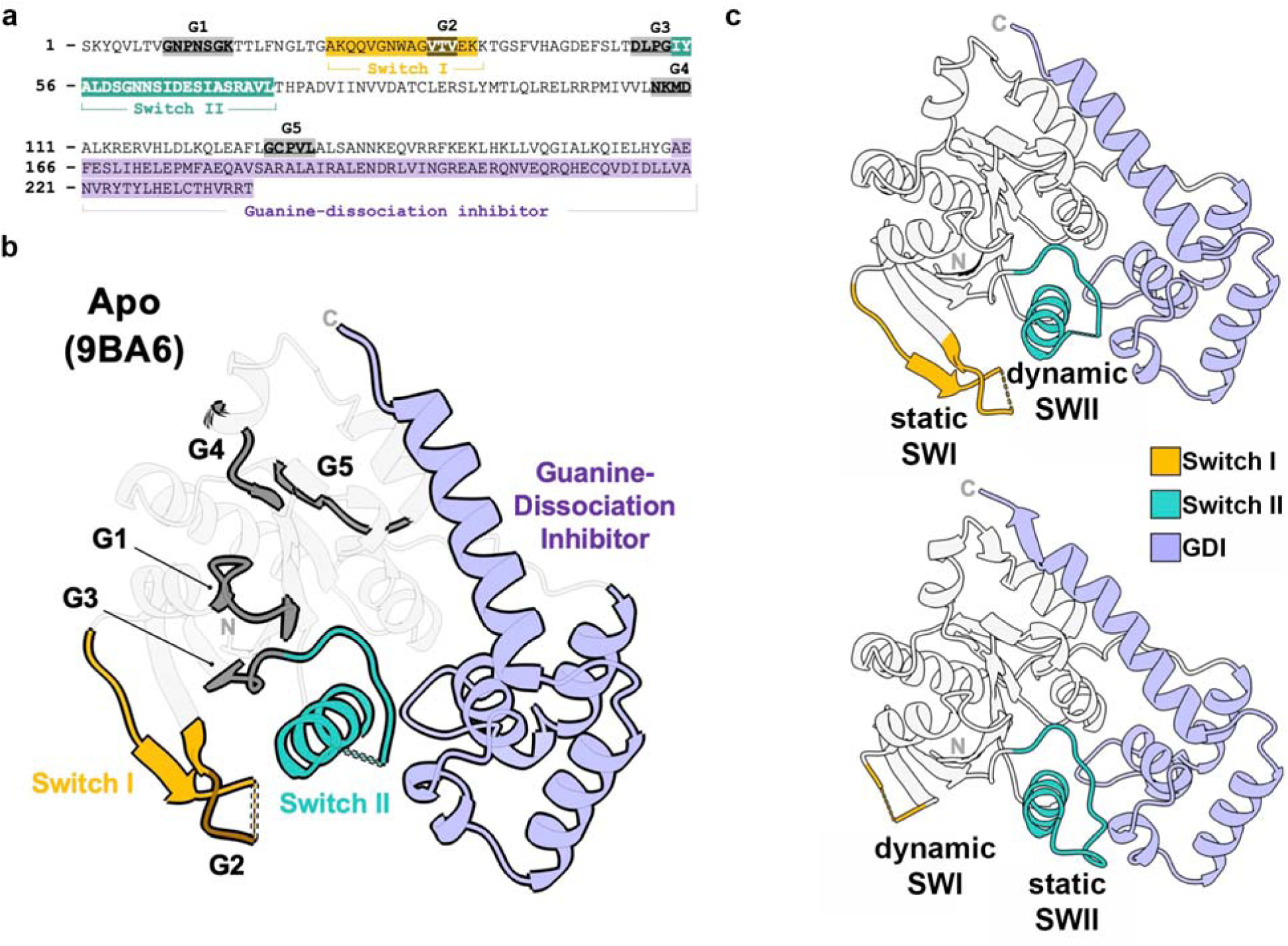
The apo structure of *Vc*NFeoB (PDB ID 9BA6). **a**. Sequence of the *Vc*NFeoB domain crystallized in this work with the various G motifs highlighted in gray and brown, Switch I (SWI) highlighted in yellow, Switch II (SWII) highlighted in teal, and the guanine-dissociation inhibitor (GDI) sub-domain highlighted in purple. **b**. The apo *Vc*NFeoB structure with the same sub-domains in panel **a** colored on the three-dimensional structure. **c**. Comparison of the apo *Vc*NFeoB protomer displaying a static SWI and a dynamic SWII (*top panel*) to the apo *Vc*NFeoB protomer displaying a dynamic SWI and a static SWII (*bottom panel*). In all cases, the labels ‘N’ and ‘C’ represent the locations of the N- and C-termini, respectively.

**Table 2.** Composition and dynamics of the asymmetric units contained in the X-ray crystal structures of apo WT*Vc*NFeoB (PDB ID 9BA6), WT*Vc*NFeoB + GDP (PDB ID 9D8B), WT*Vc*NFeoB + GMP-PCP (PDB ID 9D8D), and WT*Vc*NFeoB + GDP +AlF_3_ (PDB ID 9PSD).

| Protein crystal | Apo<br>WTVcNFeoB | WTVcNFeoB + GMP-PCP | WTVcNFeoB + GDP +AlF<sub>3</sub> | WTVcNFeoB + GDP |
| --- | --- | --- | --- | --- |
| **PDB ID** | 9BA6 | 9D8D | 9PSD | 9D8B |
| **Space group** | $P_{121}$ | $P_1$ | $P_1$ | $P_1$ |
| **Resolution (Å)** | 2.38 | 2.11 | 2.00 | 1.82 |
| **Copies of VcNFeoB in the ASU** | 2 | 4 | 4 | 4 |
| **Type & number of nucleotide(s) present** | - | 4 GMP-PCP (chains A, B, C, D) | 4 GDP (chains A, B, C, D) | 4 GDP (chains A, B, C, D) |
| **Mg<sup>2+</sup> present?** | No | Yes, in 2 protomers of the ASU (chains C, D) | Yes, in 2 protomers of the ASU (chains B, C) | No |
| **AlF<sub>3</sub> (transition state) present?** | No | No | Yes, in 1 protomers of the ASU (chain B) | No |
| **SWI ordered?** | Yes, partially in 1 protomer of the ASU (chain A) | Yes, in 2 protomers of the ASU (chains A, B) | Yes, in 3 protomers of the ASU (chain A, B*, D) | Yes, in 2 protomers of the ASU (chains A, D) |
| **SWI disordered?** | Yes, in 1 protomer of the ASU (chain B) | Yes, in 2 protomers of the ASU (chains C, D) | Yes, in 1 protomers of the ASU (chain C) | Yes, in 2 protomers of the ASU (chains B, C) |
| **SWII ordered?** | Yes, in 1 protomer of the ASU (chain B) | Yes, in 1 protomer of the ASU (chains A) | Yes, in 1 protomer of the ASU (chain D) | No |
| **SWII disordered?** | Yes, partially in 1 protomer of the ASU (chain A) | Yes, partially in 3 protomers of the ASU (chains B, C, D) | Yes, partially in 3 protomers of the ASU (chains A, B, C) | Yes, partially in 4 protomers of the ASU (chains A, B, C, D) |
| *Switch 1 in novel conformer of random coil pointing *away* from nucleotide-binding site |  |  |  |  |

### Mg^2+^ binding activates Switch I and Switch II in the GMP-PCP-bound form of VcNFeoB

The next step in the nucleotide hydrolytic cycle is binding of an NTP, and we have crystallized *Vc*NFeoB in the presence of GMP-PCP (a GTP analog) both with and without Mg^2+^ in the same ASU (Fig. 3; Table 1; PDB ID 9D8D; 2.11 Å resolution). The crystallization of *Vc*NFeoB in the presence of a GTP analog but the absence of Mg^2+^ is noteworthy, as all deposited NFeoB structures in the presence of a GTP analog contain at least one Mg^2+^ coordinated between the β- and γ-phosphates. Yet, despite co-crystallization of *Vc*NFeoB in the presence of excess GMP-PCP and Mg^2+^, we observe clear and unambiguous density for only the GTP analog in 2 molecules of the ASU (chains A and B) whereas we observe clear and unambiguous density for both the GTP analog and Mg^2+^ in the other 2 molecules of the ASU (chains C and D) (Fig. S4). In the absence of Mg^2+^, GMP-PCP is observed to be bound as expected in the G-protein like domain of chains A and B, with residues comprising the G1 through G3 motifs contributing to binding of the phosphate regions, while residues comprising the G4 and G5 motifs cradle the guanine nucleobase (Fig. S4). In the presence of Mg^2+^, GMP-PCP is bound in the identical nucleotide-binding pocket of chains C and D with only minor changes observed locally, such as the presence of a typical hexacoordinate Mg^2+^ between the β- and γ-phosphates and Thr16 of the G2 motif coordinated to the Mg^2+^ *trans* to the bound γ-phosphate oxygen (Fig. S4). In all chains, binding of the GTP analog does not elicit a gross overall conformational change compared to the apo form (core NFeoB Cα RMSD average of 0.74 Å across all four protomers in the ASU); however, subtle but important changes occur in Switch I and Switch II within the GMP-PCP-bound protomers of the ASU depending on cation binding.

**Figure 3.**
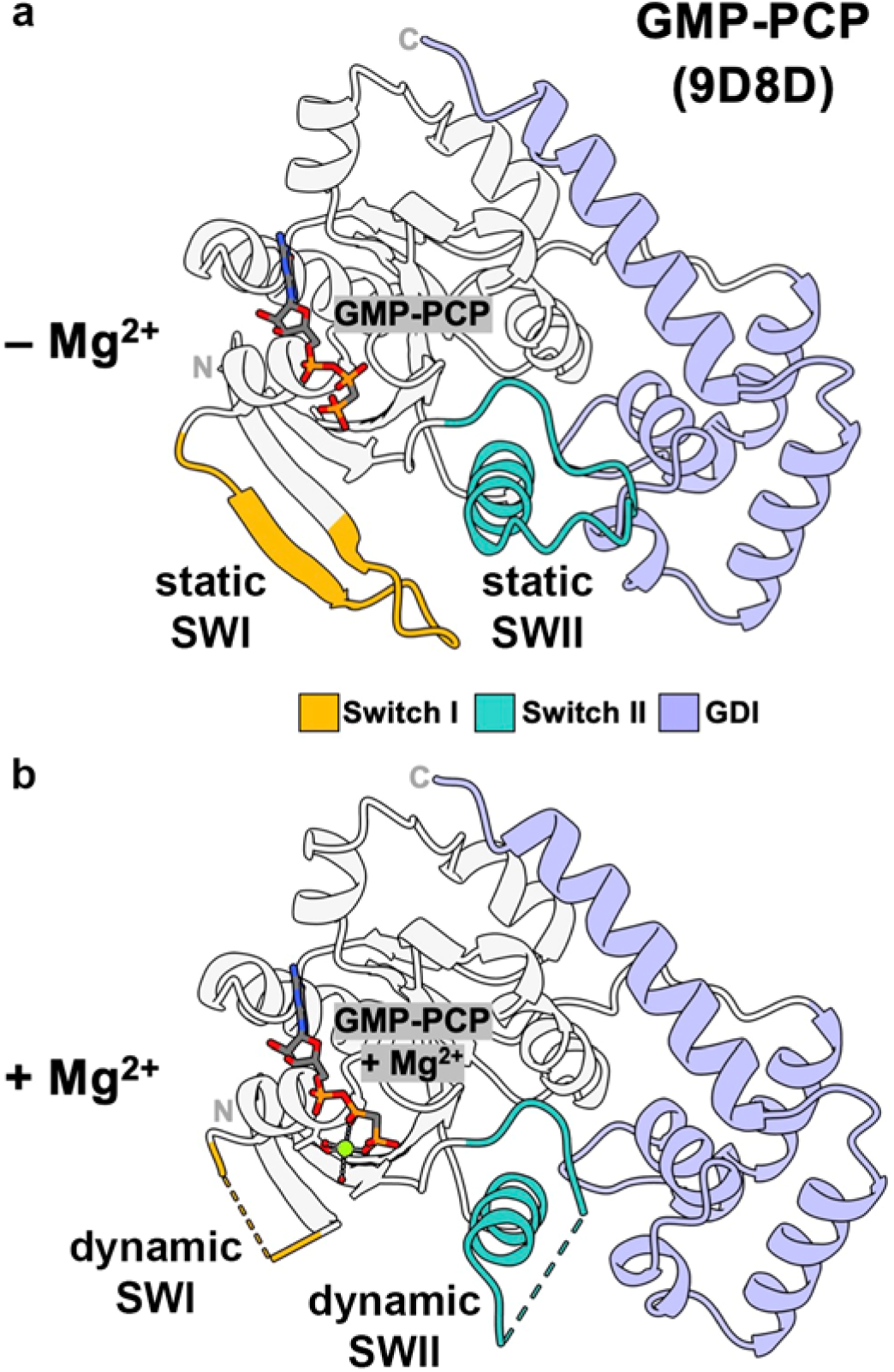
The structure of *Vc*NFeoB bound to a GTP analog (GMP-PCP; PDB ID 9D8D) in the presence (**a**) and the absence (**b**) of Mg^2+^. Switch I (SWI) is highlighted in yellow, Switch II (SWII) is highlighted in teal, and the guanine-dissociation inhibitor (GDI) sub-domain is highlighted in purple. In all cases, the labels ‘N’ and ‘C’ represent the locations of the N- and C-termini, respectively.

In our *Vc*NFeoB structure bound to GMP-PCP, coordination of Mg^2+^ between the β- and γ-phosphates of the GTP analog appears to mobilize the dynamics of Switch I and Switch II (Fig. 3; Fig. S3). For example, when Mg^2+^ is absent but GMP-PCP is present, Switch I is completely ordered in both chains (A and B), while Switch II is completely ordered in one chain (A) and partially ordered in another chain (B) (Table 2; Fig. S3). These results suggest that, in the absence of Mg^2+^, binding of the GTP analog appears to elicit a stabilizing effect on the Switch I and II regions, especially when compared to the apo form of *Vc*NFeoB (Fig. 2), likely to relay the presence of the triphosphate nucleotide to the rest of the protein. In a marked difference, when Mg^2+^ and GMP-PCP are both present (chains C and D), Switch I becomes completely disordered in both chains, accompanied by a partial disorder in Switch II generally extending from Asp63 to Ser70 (Fig. 3; Table 2; Fig. S3), similar in extent to the chain B apo form of *Vc*NFeoB (Fig. 2). The behavior of Switch I and Switch II of *Vc*NFeoB bound to Mg^2+^-coordinated GMP-PCP is comparable to what is observed in the structure of *Kp*NFeoB bound to Mg^2+^-coordinated GMP-PNP^46^ but is in contrast with the Mg^2+^-coordinated GMP-PNP structures of thermophilic NFeoBs (such as *St*NFeoB) that have captured Switch I stabilized and coordinated to Mg^2+^ via the highly conserved G2 Thr residue^51^. Importantly, the structure of Mg^2+^-free GMP-PCP bound to *Vc*NFeoB has not been previously captured and, when combined with our Mg^2+^-coordinated GMP-PCP structure, suggests a key role for Mg^2+^ in mobilizing Switch I for initiation of NTP hydrolysis.

### The transition-state analog structure of VcNFeoB reveals an open Switch I conformation

To probe the mechanism of NTP phosphate bond cleavage, the next step in the nucleotide hydrolytic cycle, we have crystallized *Vc*NFeoB in the presence of GDP, Mg^2+^, and AlF_3_, resulting in a structure that provides multiple snapshots of this step within the ASU (Fig. 4; Table 1; PDB ID 9PSD; 2.00 Å resolution). We initially attempted to capture the transition state structure of *Vc*NFeoB by co-crystallizing the protein at 20 °C with GDP, Mg^2+^, NaF, and AlCl_3_, an approach that was previously successful for *St*NFeoB, the only known transition-state analog-bound structure of NFeoB^40^. Despite this attempt, we were only able to generate crystals with GDP bound to the protein, suggesting that the transition-state analog of *Vc*NFeoB is not stable at this temperature. We were ultimately only able to capture the transition-state analog bound to *Vc*NFeoB in crystals that were generated at 4 °C, although only one molecule in the ASU contained the analog (Table 2). In chain B of the ASU, we observe clear and unambiguous density for GDP bound to Mg^2+^ and AlF_3_ in *Vc*NFeoB for the first time (Fig. S5). In contrast to the transition-state analog in chain B, in chain C of the ASU, we observe clear and unambiguous density for GDP bound only to Mg^2+^ at the β-phosphate (Fig. S5), an uncommonly captured NFeoB state^52^ that represents phosphate release prior to Mg^2+^ release. The remaining two chains of the ASU (A and D) are clearly bound to GDP without Mg^2+^ and without AlF_3_ (Fig. S5). As observed in our other crystal structures (*vide supra*), there is minimal change in the gross overall conformations of the protomers in the structure with the transition-state analog compared to the GMP-PCP-bound structure (core NFeoB Cα RMSD average of 0.62 Å across all four protomers in the ASU) and the apo structure (core NFeoB Cα RMSD average of 0.77 Å across all four protomers in the ASU). However, there are important and noteworthy changes in the Switch I, Switch II, and GDI regions, especially for the protomer bound to GDP-Mg^2+^-AlF_3_.

**Figure 4.**
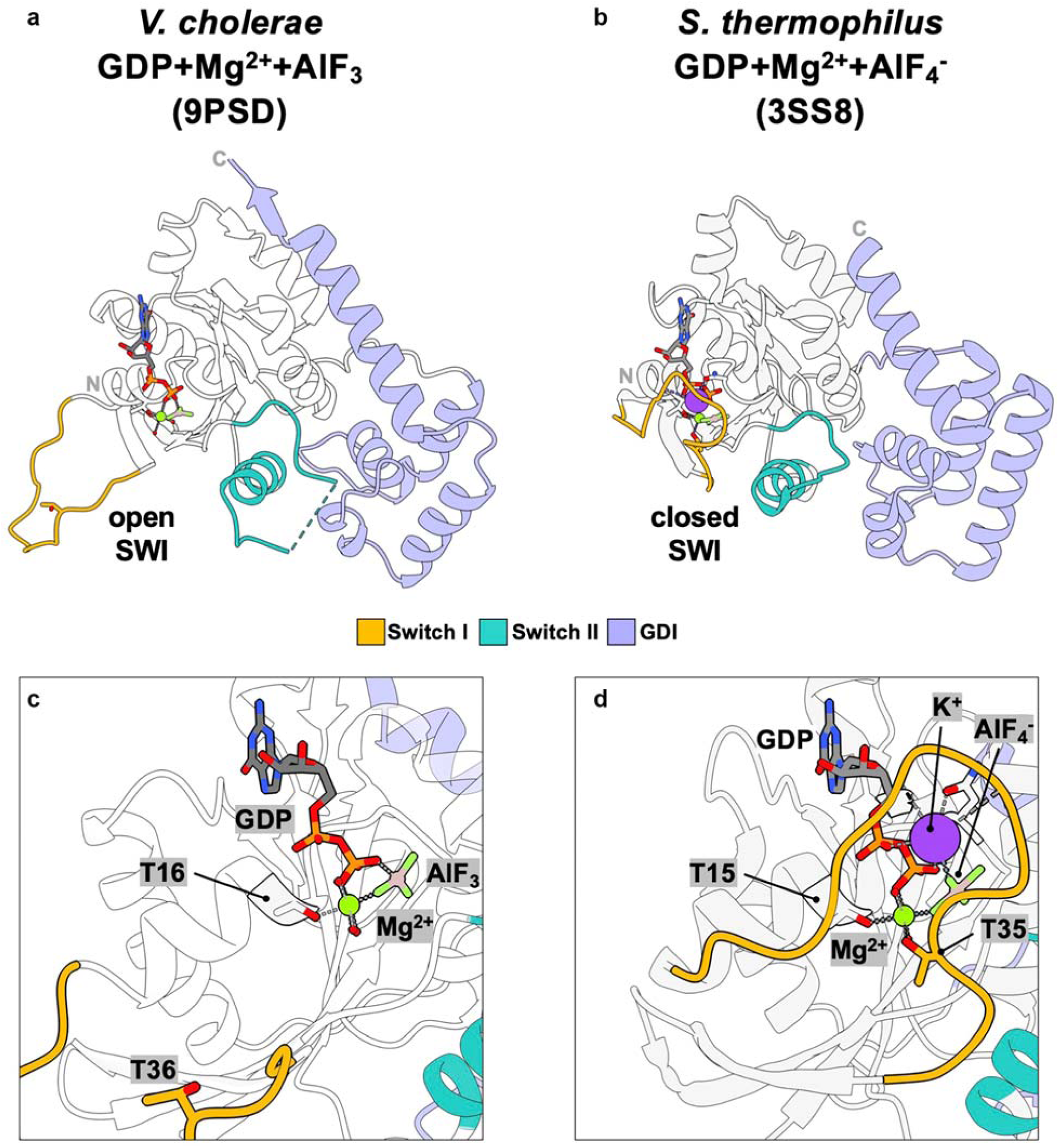
The structure of *Vc*NFeoB bound to a transition-state analog (PDB ID 9PSD) reveals an open Switch I conformation. **a**. Overall structure of *Vc*NFeoB bound to GDP-Mg^2+^-AlF_3_ compared to **b** the structure of *St*NFeoB bound to GDP-Mg^2+^-AlF_4_^-^ (PDB ID 3SS8) with a closed Switch I conformation. Panels **c** and **d** represent close-up views of the nucleotide-binding pockets of panels **a** and **b**, respectively, and highlighted are the G1 Thr residues that coordinate Mg^2+^, the SWI G2 Thr residues that are important for nucleotide hydrolysis, and the GDP molecule bound to the transition-state analogs. Switch I (SWI) is colored in yellow, Switch II (SWII) is colored in teal, and the guanine-dissociation inhibitor (GDI) sub-domain is highlighted in purple. In all cases, the labels ‘N’ and ‘C’ represent the locations of the N- and C-termini,

Within the crystals generated in the presence of the transition-state analog, two captured snapshots of *Vc*NFeoB display key structural elements that have not been previously observed. First, and most strikingly, is the observation that Switch I adopts an open conformation in the GDP-Mg^2+^-AlF_3_-bound form of *Vc*NFeoB (Fig. 4a; chain B in PDB ID 9PSD). The thermophilic *St*NFeoB is the only form of NFeoB that has been structurally characterized in the presence of a transition-state analog (GDP-Mg^2+^-AlF_4_ ^-^)^40^, but in this structure Switch I was stabilized by coordination between the G2 Thr35 (*St*NFeoB numbering) and Mg^2+^, resulting in a covering of the hydrolyzed nucleotide state that would prohibit/occlude phosphate loss (Fig. 4b). In strong contrast, our new structure of *Vc*NFeoB bound to GDP-Mg^2+^-AlF_3_ clearly reveals a conformation in which Switch I is in its open state (Fig. 4a; Figs. S3 and S5), due to a lack of coordination at the nucleotide Mg^2+^. Specifically, in the *Vc*NFeoB transition state structure, the position that is occupied by the Mg^2+^-coordinated G2 Thr35 in *St*NFeoB is instead occupied by water (W218), while the hydroxyl group of the G2 Thr36 (*Vc*NFeoB numbering) in Switch I is located *ca.* 16 Å away from the bound Mg^2+^ (Fig. S5). Moreover, Switch II remains partially ordered in the structure bound to GDP-Mg^2+^-AlF_3_, akin to the *Vc*NFeoB GMP-PCP-bound structure. Thus, in the *Vc*NFeoB transition state structure, Switch I points away from the nucleotide-binding pocket adopting an unoccluded state to allow for phosphate release. Also noteworthy is the capture of *Vc*NFeoB in the GDP-Mg^2+^-bound form (Fig. 5a; chain C in PDB ID 9PSD). NFeoB bound to GDP-Mg^2+^ has only been previously observed once in a thermophilic structure (*Thermatoga maritima* NFeoB)^52^ and was captured due to protomer-protomer crystal contacts that sandwiched the Mg^2+^ ion between two GDP molecules at the nucleotide-binding site. Here in our *Vc*NFeoB transition-state analog structure, we were able to capture the GDP-Mg^2+^-bound structure without the aid of crystal contacts (Figs. S5, S6). Like in our *Vc*NFeoB GMP-PCP-Mg^2+^-bound structure, the presence of Mg^2+^ bound to GDP and the protein mobilizes both Switch I and Switch II that are completely and partially disordered, respectively (Table 2; Fig. S3). Interestingly, in this protomer (and only in this protomer) we observe the highest thermal factors and partial disorder within the globular “head” of the GDI domain despite this portion of the polypeptide being located near the protomer-protomer interface of the ASU (Fig. S6), which would be expected to otherwise stabilize this region. The *Vc*NFeoB structure bound to GDP-Mg^2+^ suggests that, after nucleotide hydrolysis in NFeoB, loss of phosphate may occur prior to loss of Mg^2+^, and this information may be communicated through the protein by high dynamics in the GDI domain. When taken together, these results shed light on the nucleotide hydrolysis mechanism catalyzed by *Vc*NFeoB.

**Figure 5.**
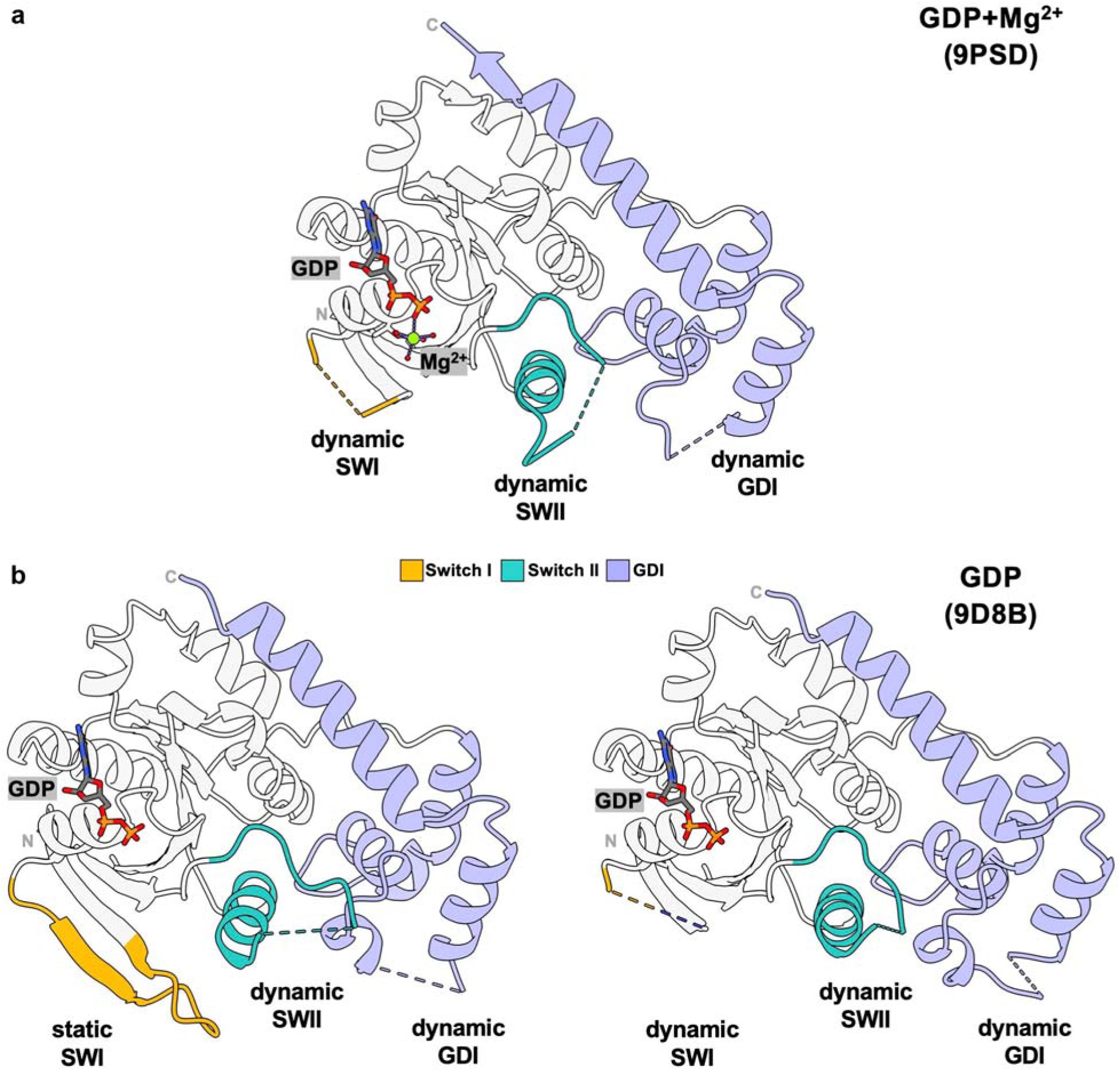
The structure of *Vc*NFeoB bound to GDP with Mg^2+^ (PDB ID 9PSD) and without Mg^2+^ (PDB ID 9D8B). **a**. The structure of *Vc*NFeoB bound to GDP and Mg^2+^ displays dynamic Switch I (SWI; yellow), Switch II (SWII; teal), and guanine-dissociation inhibitor (GDI; purple) sub-domains, characterized by multiple regions of disorder. **b**. Comparison of Mg^2+^-free, GDP-bound *Vc*NFeoB displaying a static SWI region (*left panel*) to that of Mg^2+^-free, GDP-bound *Vc*NFeoB with a dynamic SWI region (*right panel*). In both structures, SWII and the head of the GDI sub-domain display regions of disorder. In all cases, the labels ‘N’ and ‘C’ represent the locations of the N- and C-termini, respectively.

### GDP binding to VcNFeoB in the absence of Mg^2+^ recapitulates apo Switch I dynamics

To complete the nucleotide hydrolytic cycle, we then crystallized *Vc*NFeoB in the presence of only GDP (Fig. 5b; Table 1; PDB ID 9D8B; 1.82 Å resolution). While we were previously able to crystallize this nucleotide-bound form of *Vc*NFeoB, our prior (His)_6_-tagged structure was only resolved to *ca*. 4.2 Å, limiting our ability to comment confidently on the positions of GDP, Switch I, Switch II, and the GDI domain.^31^ However, by utilizing a cleaved, (His)_6_-SUMO fusion of *Vc*NFeoB in this work, we significantly improved the quality and resolution of our derived crystals, and we observe clear and unambiguous density for GDP (and only GDP) bound in the nucleotide-binding pocket of all four protomers comprising the ASU (Fig. S7; Table 2). The overall structure of GDP-bound *Vc*NFeoB generally resembles our prior structure^31^, and there is minimal change in the gross overall conformations of the *Vc*NFeoB protomers in the GDP-bound structure compared to the GMP-PCP-bound structure (core NFeoB Cα RMSD average of 0.65 Å across all four protomers in the ASU) and compared to the apo structure (core NFeoB Cα RMSD average of 0.75 Å across all four protomers in the ASU). Interestingly, in our *Vc*NFeoB GDP-bound structure, we observe a pattern of Switch I order/disorder that appears to mirror our apo structure: in two copies of the ASU (chains A and D) Switch I is entirely ordered, while in two copies of the ASU (chains B and C) Switch I is nearly entirely disordered (generally from residues Gly24 through Lys40) (Table 2; Fig. S3). In contrast to the apo structure, and despite the high resolution of the data (1.82 Å), in the GDP-bound *Vc*NFeoB structure we observe partial disorder in Switch II in every protomer of the ASU, generally within the Ser- and Asn-rich loop that spans between Asp63 and Ile71 (Table 2; Fig. S3). This observation contrasts with most (but not all) GDP-bound NFeoB structures that appear to have both Switch I and Switch II resolved, and we believe these data point to a return of the catalytic cycle to apo-like dynamics in the GDP-bound form, at least in the Switch I region.

### The presence of Mg^2+^ impacts the ability of VcNFeoB to bind nucleotides

Finally, as our structural data point to a critical role for Mg^2+^ binding in activating both Switch I and Switch II of the *Vc*NFeoB domain, we sought to use isothermal titration calorimetry (ITC) to discern whether Mg^2+^ would impact nucleotide binding to the protein in solution (Fig. 6). As an initial check, we first determined the binding thermodynamics of GDP to our SUMO-cleaved *Vc*NFeoB in the absence of Mg^2+^ that was fitted to a single binding site with a K_d_ of 2.91 μM ± 0.34 μM (Fig. 6a), nearly identical to our prior results with the (His)_6_-tagged protein^31^ that displayed a single binding site for GDP with a K_d_ of *ca*. 3 μM in the absence of Mg^2+^. Interestingly, the presence of Mg^2+^ modestly destabilized the binding of GDP to *Vc*NFeoB by increasing the K_d_ to 16.17 μM ± 2.34 μM (Fig. 6b). As we observed the most noteworthy changes in *Vc*NFeoB structure when bound to GMP-PCP and Mg^2+^, we then determined these binding thermodynamics via ITC. As expected, in the absence of Mg^2+^, GMP-PCP bound to *Vc*NFeoB weaker than GDP with a single binding site fitted to a K_d_ of 47.10 μM ± 7.99 μM (Fig. 6c), comparable to the (His)_6_-tagged protein that displayed a fitted K_d_ for GMP-PNP of *ca*. 121 μM^31^ in the absence of Mg^2+^. Remarkably, however, we failed to observe any measurable binding of GMP-PCP to *Vc*NFeoB in the presence of Mg^2+^, even at high solution stoichiometries (> 10 mol. eq.) of the nucleotide analog relative to protein (Fig. 6d). These data contrast sharply with prior stopped-flow experiments utilizing fluorescently-tagged nucleotide analogs that measured *k_on_* and *k_off_* rates or intrinsic Trp fluorescence in order to compute the K_d_s of various GTP analogs.^40, 49, 51, 53–54^ However, these approaches did not measure K_d_ directly, not all measurements were made in the presence of Mg^2+^, and it is possible that *Vc*NFeoB may display this unique behavior due to its documented nucleotide promiscuity^30–31, 33^. Thus, our ITC data suggest a mechanism for *Vc*NFeoB-catalyzed GTP hydrolysis in which GTP binds to the nucleotide-binding pocket *prior* to Mg^2+^ binding; moreover, once nucleotide is hydrolyzed, GDP binds tightest to *Vc*NFeoB only *after* phosphate and Mg^2+^ have been released. When considered all together, these structural and biophysical data paint a comprehensive picture of *Vc*NFeoB-catalyzed GTP hydrolysis, as discussed below.

**Figure 6.**
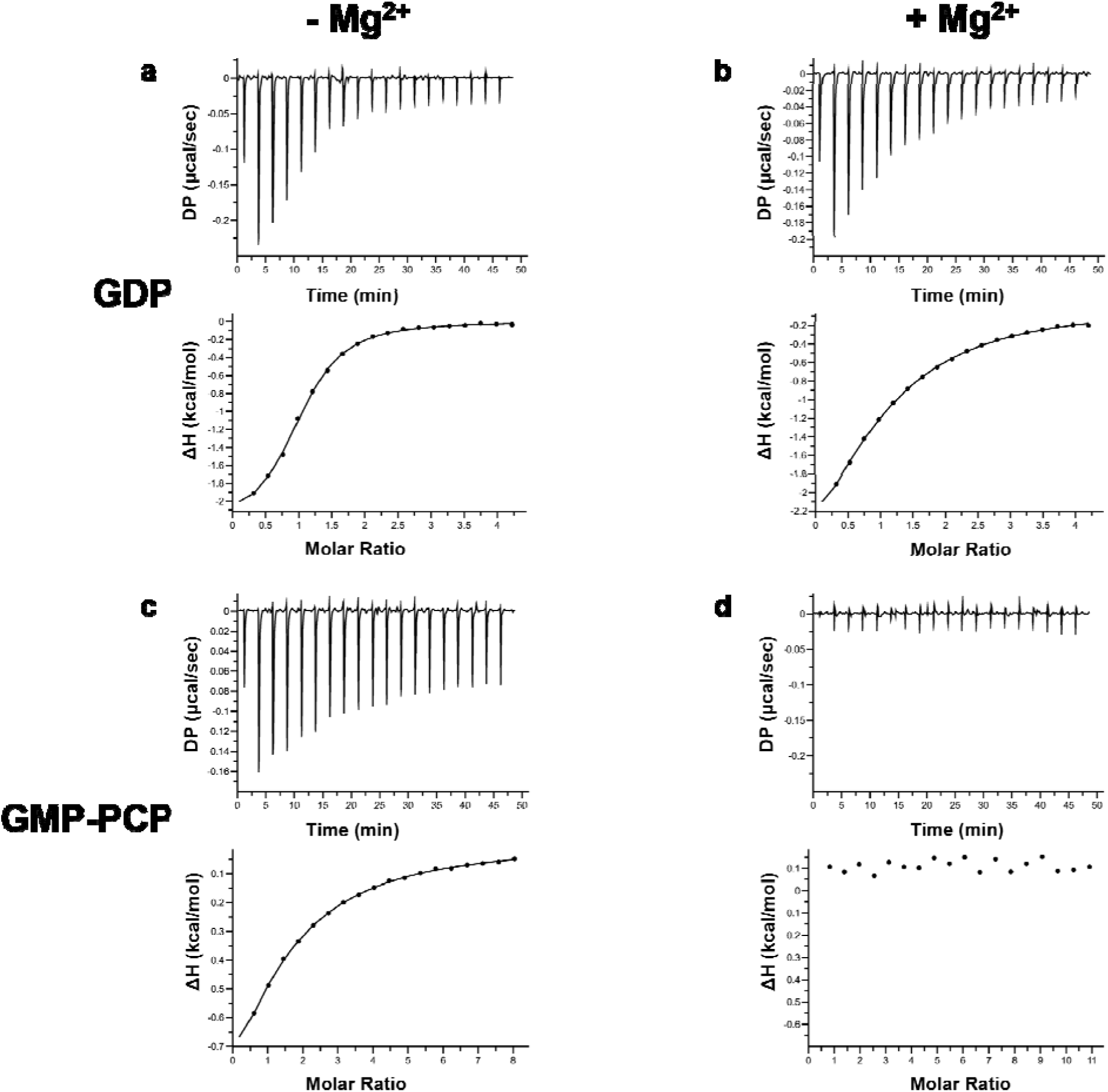
The presence of Mg^2+^ affects the ability of *Vc*NFeoB to bind nucleotides. Representative ITC thermograms (*top*) and ΔH vs. molar ratio traces (*bottom*) of *Vc*NFeoB in the absence of Mg^2+^ (*left*) and the presence of Mg^2+^ (*right*) titrated with either GDP (**a**, **b**) or GMP-PCP (**c**, **d**). All datasets have been corrected for nucleotide dilution into buffer in the absence of protein. The GDP K_d_s in the absence of Mg^2+^ (2.91 μM ± 0.34 2+ μM) and the presence of Mg^2+^ (16.17 μM ± 2.34 μM) differ modestly, with both having a single binding site (N ≈ 1). As expected, GMP-PCP binds to *Vc*NFeoB at a single binding site (N ≈ 1) more weakly (K_d_ 47.10 μM ± 7.99 2+ μM) than GDP in the absence of Mg^2+^ (**c**), while GMP-PCP markedly fails to bind to *Vc*NFeoB in the presence of Mg^2+^. All values were determined in triplicate and represent the mean ± one standard deviation of the mean.

## DISCUSSION

In this study, we have determined several new X-ray crystal structures allowing us to model the full catalytic cycle of *V. cholerae* NFeoB (Fig. 7). While the NFeoB domain has been biochemically and structurally characterized from a number of bacterial species^44–47^, the complete catalytic cycle of an NFeoB from a single organism has remained elusive, especially as the transition-state analog structure has only been determined for an NFeoB from a single, thermophilic bacterium, *S. thermophilus*^40^. Moreover, as *V. cholerae* (N)FeoB is a promiscuous NTPase, rather than a strict GTPase,^27, 30–33^ we wondered whether the structural basis of NTP hydrolysis of this NFeoB would mirror or diverge from other GTP-strict NFeoBs as well as other well-known NTPases. To answer this question, we leveraged our previous approach^31^ and set out to crystallize *Vc*NFeoB with various nucleotides. Uniquely, our newly-identified conditions facilitated the crystallization of the *Vc*NFeoB domain in the *P_1_* space group containing four non-oligomeric protomers in the ASU, fortunately allowing us to observe distinct snapshots of the *Vc*NFeoB domain in uncommon or unprecedented states, ultimately providing important mechanistic insight into this specific system and to (N)FeoB function as a whole.

**Figure 7.**
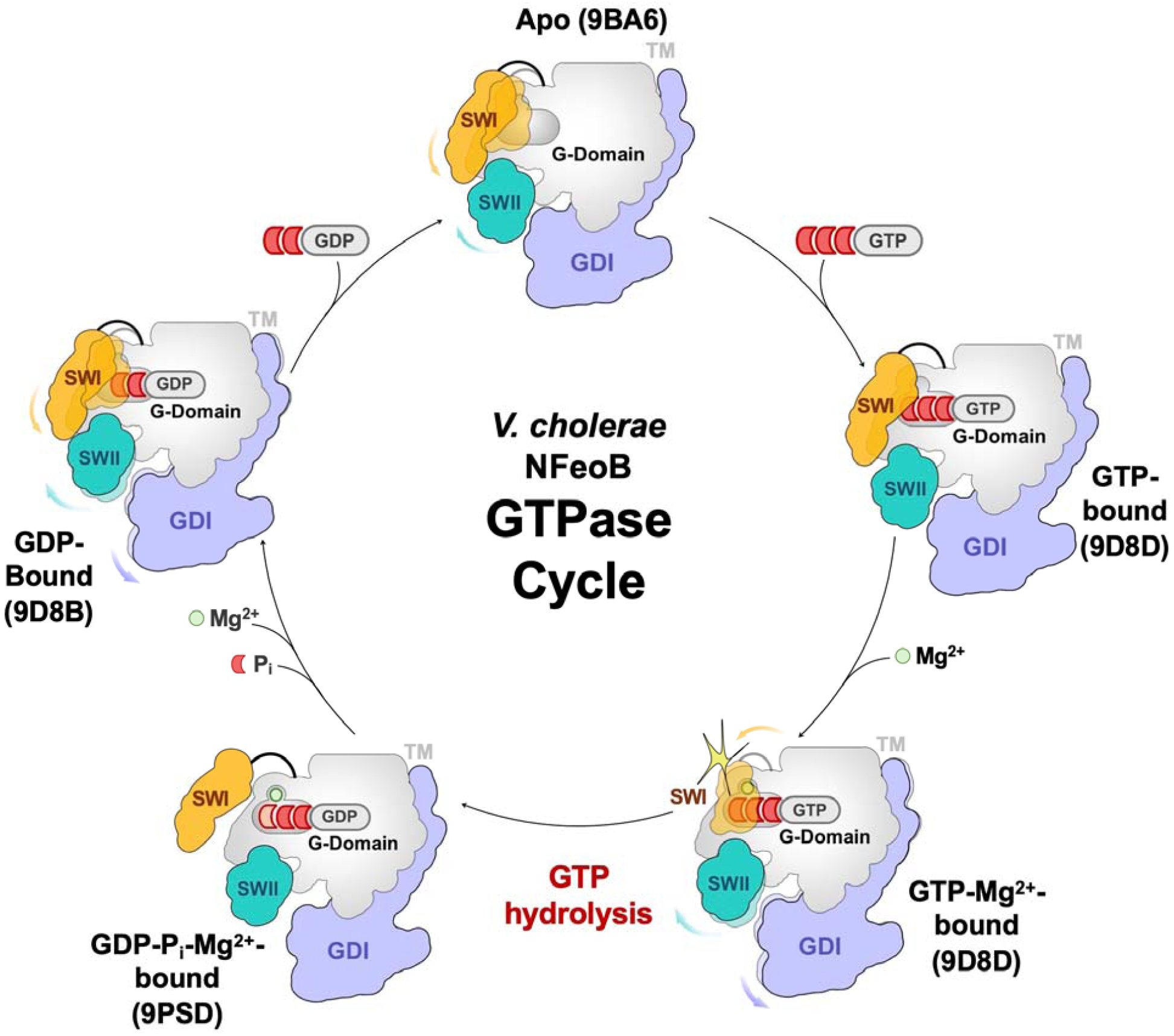
Cartoon of the GTP hydrolysis cycle of *V. cholerae* NFeoB. The apo form of *Vc*NFeoB displays dynamic Switch I (SWI; yellow) and Switch II (SWII; teal) regions prior to nucleotide binding. GTP binds to the apo state first, stabilizing SWII and SWII, after which Mg^2+^ binds to the GTP-bound form of *Vc*NFeoB, mobilizing SWI, SWII, and the guanine-dissociation inhibitor (GDI; purple) sub-domains. Upon GTP hydrolysis, SWI adopts an open conformation to allow phosphate (P_i_) release prior to Mg^2+^ release, after which the G-domain tightens its binding to GDP while the rest of *Vc*NFeoB exhibits mobility in SWI, SWII, and the GDI sub-domains. The factors affecting GDP loss and GTP rebinding remain unclear. In this cartoon, ‘TM’ represents the connection from the GDI region to the transmembrane domain of FeoB.

Our determined structures, combined with our biophysical characterization, reveal insight into conformational and dynamical changes that occur in the Switch I, Switch II, and GDI sub-domains that appear to be both important and noteworthy for the *Vc*NFeoB catalytic cycle. For example, in our apo *Vc*NFeoB structure, we observe different poses in both the Switch I and Switch II regions (static and resolved in some cases, dynamic and unresolved in others) that suggest these critical communication elements within the NFeoB domain are highly mobile in the absence of nucleotide (Fig. 7). Binding of a GTP analog (GMP-PCP) appears to stabilize these domains away from the nucleotide-binding site, at least in the absence of Mg^2+^. Once Mg^2+^ binds, Switch I and Switch II are mobilized, which is especially important for the coordination of the Switch I G2 Thr36 to Mg^2+^ in order to initiate nucleotide hydrolysis (Fig. 7). These results suggest an order of operations in which NTP binds prior to Mg^2+^ binding in *Vc*NFeoB, and our ITC data reveal that a GTP analog only binds to *Vc*NFeoB in the absence of Mg^2+^, in contrast with prior fluorescence results^40, 49, 51, 53–54^, although it is not entirely clear how many of these previous measurements were made in the presence or absence of Mg^2+^. In another contrast, our transition-state structure reveals that *Vc*NFeoB adopts an open Switch I conformation under these crystallization conditions post nucleotide hydrolysis, which would expose the nucleotide-binding pocket for phosphate release but is markedly different than the closed/occluded transition-state analog-bound structure of the thermophilic *St*NFeoB^40^. In the same crystal as the transition-state analog, we also observe the GDP-Mg^2+^-bound form of *Vc*NFeoB, a rarely-captured state of the protein representing post phosphate release but with the retention of Mg^2+^. This structure, which also displays a highly mobile, partially disordered GDI domain, implies that phosphate may exit NFeoB prior to Mg^2+^ loss (Fig. 7), suggesting that both the entrance and exit of Mg^2+^ into the nucleotide-binding pocket are highly ordered. Finally, once both phosphate and Mg^2+^ vacate the nucleotide-binding pocket, the protein tightens its grip on the hydrolyzed NDP while Switch I begins to resemble the apo state and both Switch II and the GDI domain remain partially mobile and disordered (Fig. 7). When taken together, these data provide a structural basis of the nucleotide hydrolytic cycle catalyzed by *Vc*NFeoB.

However, it is important to remember that FeoB does not function on its own and has partner proteins such as FeoA (highly conserved) and FeoC (poorly conserved) that may target key regions identified in these structures to control function.^6, 21, 25–26^ Interestingly, previous research has shown that FeoA and FeoC are known to bind to the NFeoB domain under different conditions, and their binding sites appear to be mutually overlapping. Regarding FeoA, *in vitro* studies of the naturally-occurring *Bacteroides fragilis* FeoA-NFeoB fusion demonstrated that FeoA interacts with NFeoB in a nucleotide-dependent manner, with the tightest interaction being in the presence of a GTP analog but in the absence of Mg^2+^.^42^ In this system, the FeoA binding site on NFeoB included various regions within the Switch I, Switch II, and GDI domains.^42^

Importantly, the presence of FeoA-NFeoB interactions stabilized and prevented nucleotide hydrolysis^42^, likely due to FeoA blocking Switch I from being mobilized for nucleotide hydrolysis. Additionally, *V. cholerae* studies have shown that *in vivo* mutations of the G1 and Switch II NFeoB regions as well as mutations in FeoA itself prevent Feo complex formation and function.^35^ Regarding FeoC, an *in vitro* co-crystal structure of *K. pnuemoniae* FeoC/NFeoB revealed binding of *Kp*FeoC via its helix-turn-helix (HTH) motif to the *Kp*NFeoB Switch II and GDI sub-domains^55^, similar to the locations along NFeoB that are implicated in complex formation in *V. cholerae*.^35^ Consistent with these results, we previously determined the NMR structure of *Vc*FeoC and used *in vitro* NMR titrations of *Vc*FeoC into intact, full-length *Vc*FeoB solubilized in detergent to show that both the HTH and the winged loop regions of *Vc*FeoC make contact with *Vc*FeoB, and modeling suggested these protein-protein interactions occur at the Switch I-Switch II-GDI interface in the cytosolic *Vc*NFeoB domain.^36^ When combined, these data point to FeoA and FeoC having mutually overlapping binding sites on NFeoB that engage the Switch I, Switch II, and the GDI sub-domains, all critical for nucleotide hydrolysis and for relaying changes in nucleotide status to the rest of FeoB. It is unlikely that FeoA and FeoC elicit the same effect on NFeoB when they bind to these sites, but more work is needed to define the effects of these intermolecular interactions and to test whether this protein-protein interface could be targeted to alter bacterial iron acquisition and pathogenesis.

## Acknowledgements

This work was supported by NIH-NIGMS grant R35 GM133497 (A.T.S.), NIH-NIGMS grant T32 GM158458 (C.A. and A.T.S.), and NSF grant 2207374 (K.M.). NSLS-2 is a U.S. DOE Office of Science User Facility operated under Contract No. DE-SC0012704. This publication resulted from the data collected using the beamtime obtained through NE-CAT BAG proposal # 317877. The authors of this work wish to thank Prof. Michael F. Summers (UMBC and HHMI) for access to the MicroCal Automated PEAQ-ITC instrumentation. Sequence searches utilized both database and analysis functions of the Universal Protein Resource (UniProt) Knowledgebase and Reference Clusters (http://www.uniprot.org) and the National Center for Biotechnology Information (http://www.ncbi.nlm.nih.gov/).

## Author contributions

K.M., C.M.A., M.L., and A.T.S. designed the research; K.M., C.M.A., and M.L. performed the research; K.M., C.M.A., and A.T.S. analyzed the data; and K.M., C.M.A., and A.T.S. wrote and edited the paper.

## Additional information

Supplementary information. Supplementary information accompanies this paper that includes supplementary figures S1-S7 as well as any additional experimental details, materials, and methods.

## Competing interests

The authors declare no competing interests.

## METHODS

### Materials

All codon-optimized genes used in this study were synthesized and verified by GenScript. Materials used for buffer preparation, protein expression, and protein purification were purchased from standard commercial suppliers and were used as received. GDP and GMP-PCP were purchased from MilliporeSigma and stored at -20 °C until used as received.

### Cloning, expression, and purification of *Vc*NFeoB

The expression plasmid for the SUMO-fused N-terminal NTPase domain of FeoB from *Vibrio cholerae* (Uniprot ID C3LP27) was generated as previously described.^31^ Briefly, the codon-optimized gene encoding for *Vc*NFeoB was synthesized with a synthetically added sequence for the Small Ubiquitin-like Modifier (SUMO) protein (Uniprot ID Q12306). The entire sequence was then subcloned into the pET-45b(+) plasmid between the *Pml*I and *Pac*I restriction sites, which conferred ampicillin resistance and allowed for the translation of the N-terminal (His)_6_-tagged SUMO-*Vc*NFeoB fusion when read in frame.

The expression, purification, and cleavage of (His)_6_-SUMO-*Vc*NFeoB was performed as previously described.^31^ Briefly, the plasmid containing (His)_6_-SUMO-*Vc*NFeoB was transformed into *Escherichia coli* BL21 (DE3) electrocompetent cells via electroporation, after which transformed cells were plated on Luria-Bertani (LB) plates supplemented with ampicillin (100 μg/ml, final) and incubated overnight at 30 °C. Single colonies were then used to inoculate overnight starter cultures for large-scale expression that was accomplished in 1 L baffled flasks charged with 1 L LB broth with ampicillin (100 μg/mL, final). Large-scale cultures were shaken at 37 °C, 200 RPM until the optical density of the culture at 600 nm (OD_600_) reached a value from 0.6 to 0.8. After this point, large-scale cultures were then cold-shocked at 4 °C for 2 hr before being induced by the addition of 1 mM (final) isopropyl β-D-1-thiogalactopyranoside (IPTG). Large-scale cultures were then shaken at 18 °C, 200 RPM for 16 to 18 hr, after which cells were harvested by centrifugation at 5000 x*g*, resuspended in resuspension buffer (50 mM Tris pH 8.0, 300 mM NaCl, and 10 % (v/v) glycerol) before being flash frozen on N_2(l)_ and stored at -80 °C. As needed, frozen cells were thawed, resuspended with an additional 50 mL resuspension buffer, treated with 1 mM (final) phenylmethylsulfonyl fluoride (PMSF), and sonicated at 80 % maximal amplitude with a 30 s on pulse and a 30 s rest pulse for 12 min total sonication time. The lysed cells were then clarified by spinning 163000 x*g* for 1 hour at 4 °C. The clarified supernatant was then applied to a 5 mL HisTrap HP column (Cytiva) pre-charged with NiCl_2_. The column was equilibrated with 5 column values (CVs) wash buffer (50 Mm Tris pH 8.0, 300 mM NaCl, 10 % (v/v) glycerol and 1 mM TCEP), after which the column was washed extensively prior to the protein being eluted with wash buffer containing an additional 150 mM imidazole. The eluted fractions containing crude (His)_6_-SUMO-*Vc*NFeoB were then buffer exchanged using a 50 mL HiPrep 26/10 desalting column into 50 mM Tris pH 8.0, 10 % (v/v) glycerol, 10 mM β-mercaptoethanol (BME) and then concentrated using a 10 kDa molecular weight cutoff (MWCO) spin concentrator. House-made SUMO protease Ulp1 was then added (generally at a 1:100 (mg/mg) ratio) and allowed to cleave overnight at room temperature. The next day, the solution was applied again to a 5 mL HisTrap HP column to separate the now cleaved *Vc*NFeoB from any uncleaved protein and the (His)_6_-SUMO moiety. Flow-through fractions containing cleaved *Vc*NFeoB were then concentrated via a 10 kDa MWCO spin concentrator, injected onto a 120 mL preparative Superdex 75 column, eluted isocratically into SEC buffer (25 mM Tris (pH 8.0), 100 mM NaCl, 5 % (w/v) glycerol, 1 mM TCEP), concentrated, flash-frozen on N_2(l)_, and stored at -80 °C.

### Crystallization of *Vc*NFeoB

SUMO-cleaved *Vc*NFeoB was initially thawed and diluted to *ca*. 8-10 mg/ml with SEC buffer prior to nucleotide binding and crystallization screening. All protein additives were prepared to their final concentrations (*vide infra*) in SEC buffer. To prepare the guanosine diphosphate (GDP)-bound form of *Vc*NFeoB, GDP was preincubated at 0.3 mM (final concentration) with the protein (8.3 mg/mL final concentration) for two hours at room temperature prior to tray setup. To prepare the guanosine-5′-[(β,γ)-methyleno] triphosphate (GMP-PCP)-bound form of *Vc*NFeoB, GMP-PCP was preincubated at 0.72 mM (final concentration) with MgCl_2_ (1 mM final concentration) and with protein (10 mg/mL final concentration) for two hours at room temperature prior to tray setup. To prepare the transition state-bound form of *Vc*NFeoB, GDP was preincubated at 0.30 mM (final concentration) with MgCl_2_ (10 mM final concentration), NaF (5 mM final concentration), AlCl_3_ (0.5 mM final concentration), and with protein (8.3 mg/mL final concentration) for two hours at room temperature prior to tray setup. Crystallization of all forms of *Vc*NFeoB occurred using vapor diffusion in sitting-drop format comprising 1 μL of protein mixed with 1 μL of precipitant. The GDP-bound form of *Vc*NFeoB was crystallized at 20 °C using precipitant composed of 0.2 M ammonium acetate, 26 % (w/v) PEG 4000, 0.1 M bicine, pH 8.4. The GMP-PCP-bound form of *Vc*NFeoB was crystallized at 20 °C using precipitant composed of 18 % (w/v) PEG 4000, 0.1 M bicine, pH 8.4. The transition state-bound form of *Vc*NFeoB was crystallized at 4 °C using precipitant composed of 25 % (w/v) PEG 3350, 0.1 M HEPES, pH 7.5. In all cases, formed crystals were then looped, cryoprotected (typically in the mother liquor containing an additional 20-30 % (v/v/) glycerol), and flash-frozen in N_2(l)_.

### X-ray diffraction, data reduction, and structural determination

Diffraction data were collected at Brookhaven National Laboratory beamline 17-ID-2 (FMX). Data were automatically processed using Xia2^56^ and/or AutoProc^57^. The initial phases of all datasets were determined by molecular replacement (MR) using Phenix Phaser^58^ with SUMO-cleaved apo *Vc*NFeoB (PDB ID 9BA6) and GDP-bound *Vc*NFeoB(His)_6_ (PDB ID 8VWN) as initial inputs.^31^ After an initial MR solution was identified, further model building was accomplished using Phenix AutoBuild^58^. Iterative rounds of manual model building and refinement were accomplished in Coot^59^ and Phenix Refine^58^, respectively, until model convergence and the final placement of visible solvent molecules. Ramachandran statistics and clash values were determined from the MolProbity program^60^ within the Phenix software suite. The following structures have been deposited in the Protein Data Bank (PDB): SUMO-cleaved *Vc*NFeoB + GDP in *P_1_* space group (PDB ID 9D8B); SUMO-cleaved *Vc*NFeoB + GMP-PCP in *P_1_* space group (PDB ID 9D8D); and SUMO-cleaved *Vc*NFeoB + GDP +AlF_3_ in *P_1_* space group (PDB ID 9PSD). Data collection and refinement statistics are provided for all structures in Table 1.

### Isothermal titration calorimetry

All isothermal titration calorimetry (ITC) experiments were conducted using the MicroCal Automated PEAQ-ITC (Malvern Analytical). The injection syringe was loaded with 40 μL of *ca*. 1-3 mM ligand (GDP or GMP-PCP), then titrated into a 200 μL calorimetry cell containing a final concentration of 25 μM SUMO-cleaved *Vc*NFeoB in SEC buffer (25 mM Tris pH 8.0, 100 mM NaCl, 5% (v/v) glycerol, 1 mM TCEP). For experiments involving Mg^2+^, a final concentration of 5 mM MgCl_2_ was added both to the syringe and sample cell. Thermal equilibrium was reached at 25 °C after an initial 60 s delay followed by 19x 1 μL serial injections into the cell with 150 s interval delays between injection points with high spinning. Resulting data were analyzed using the Malvern MicroCal PEAQ ITC analysis tool and fitted to a binding isotherm with a single site using the following equations:

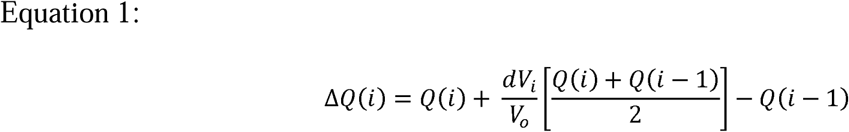

Where the heat released, ΔQ(i), from the ith injection is represented by ΔQ(i).

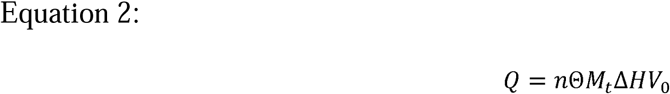

Where the total heat (Q) is related to the number of sites (n), the fractional occupation (Θ) the total free concentration of the macromolecule (*M*_t_), the molar heat of ligand binding (ΔH), and the volume determined relative to zero for the unbound species (V_0_).

## Abbreviations

CV: column volumes
Feo: ferrous iron uptake system
GDP: guanosine diphosphate
GMP-PCP: guanosine-5′-[(β,γ)-methyleno] triphosphate
GMP-PNP: guanosine-5′-[(β,γ)-imido] triphosphate
IMAC: immobilized metal affinity chromatography
IPTG: isopropyl β-D-1-thiogalactopyranoside
ITC: isothermal titration calorimetry
LB: Luria broth
NFeoB: N-terminal NTPase domain of FeoB
PMSF: phenylmethylsulfonyl fluoride
SEC: size-exclusion chromatography
ULP1: Ubiquitin-like-specific protease 1
WT: wildtype.

## Notes

### Competing Interest Statement

The authors have declared no competing interest.

