## Supporting Information for "The structure of the full catalytic cycle of *Vibrio cholerae* NFeoB"

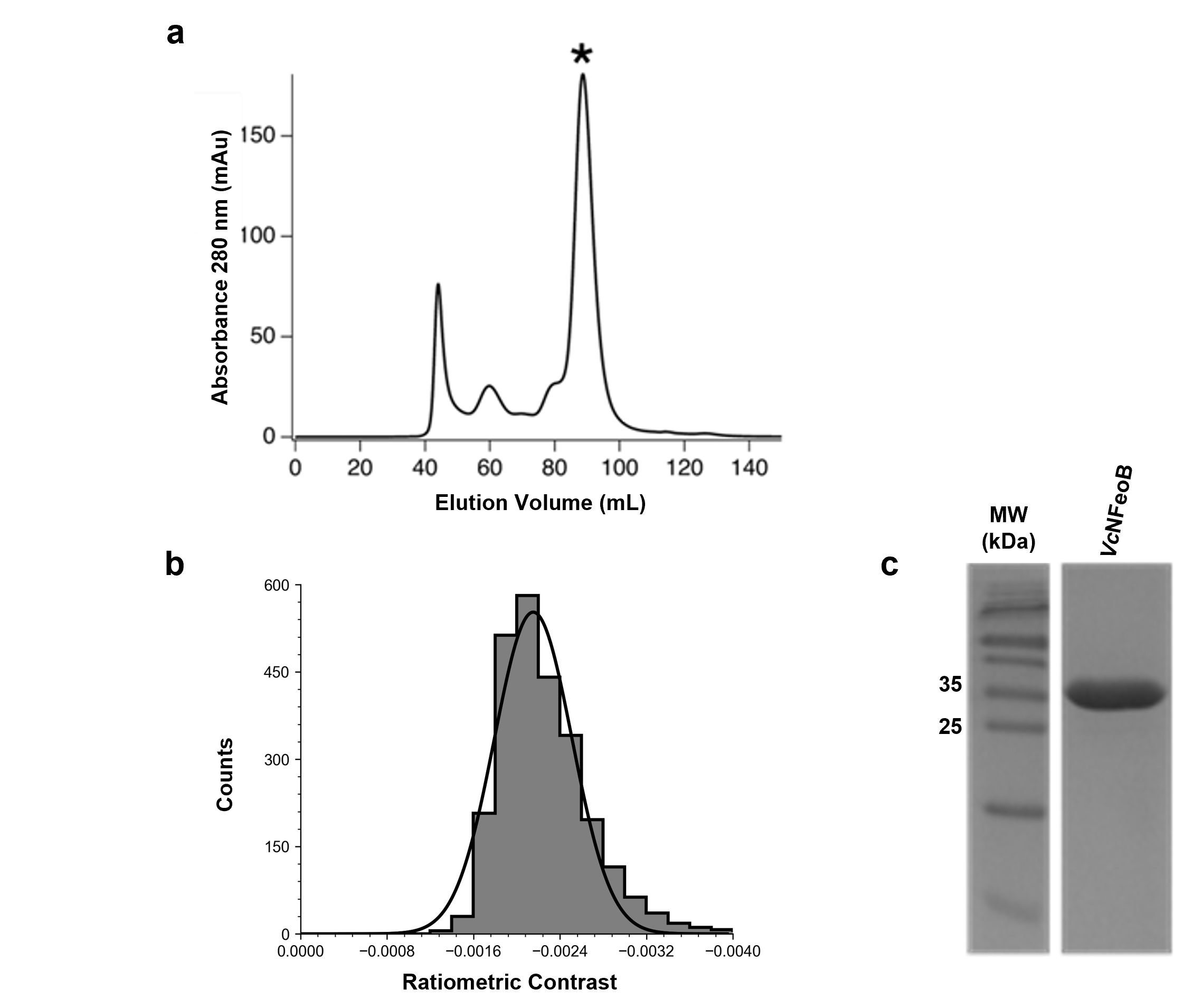

**Figure S1**. Purity and homogeneity of SUMO-cleaved *Vc*NFeoB. **a**. Preparative size-exclusion chromatogram (SEC) of *Vc*NFeoB post cleavage. The asterisk (*) represents the presence of monomeric *Vc*NFeoB. **b**. In-solution mass photometry of purified *Vc*NFeoB demonstrating its monodispersity post SEC. **c**. 15 % SDS-PAGE analysis of purified, monodisperse *Vc*NFeoB post SEC (theoretical molecular weight is 29.2 kDa).

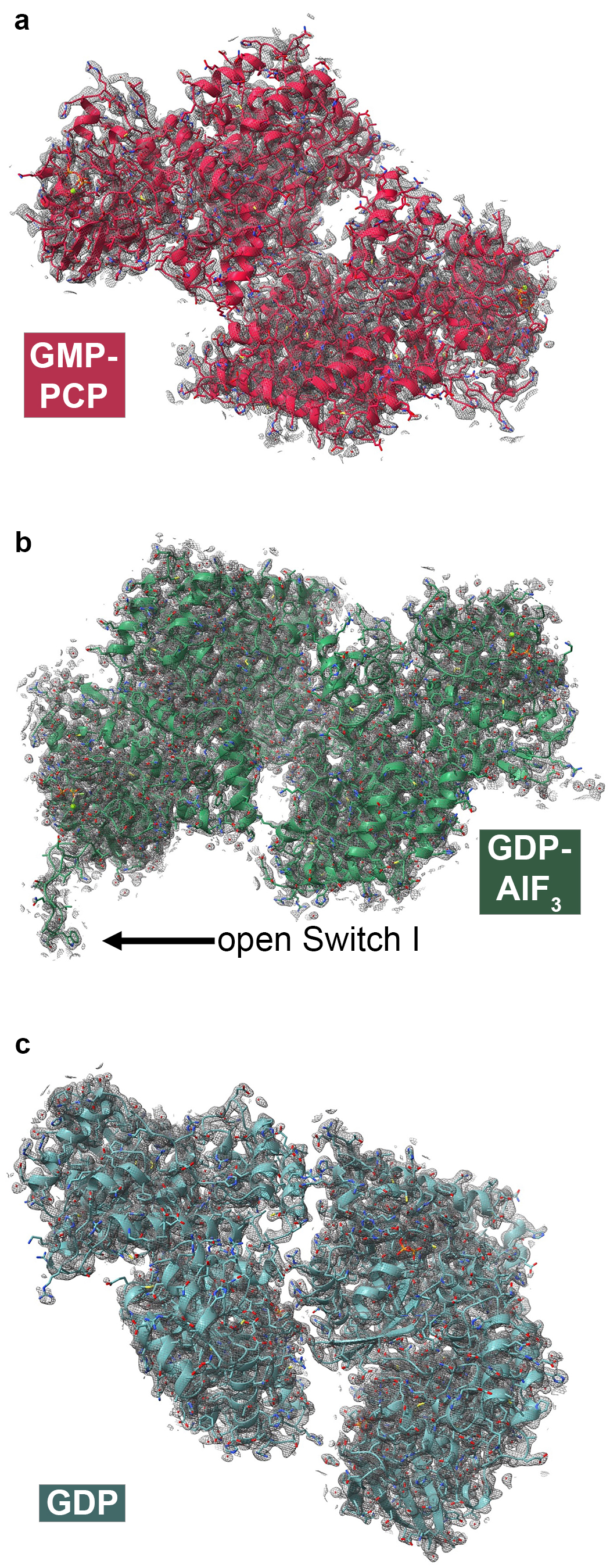

**Figure S2**. Quality of the 2*F_o_*-*F_c_* electron density contoured to 1σ (colored gray) for the entire asymmetric units (ASUs) of (**a**) the GMP-PCP-bound form of *Vc*NFeoB (red), (**b**) the GDP-AlF_3_-bound form of *Vc*NFeoB (green), and (**c**) the GDP-bound form of *Vc*NFeoB (teal).

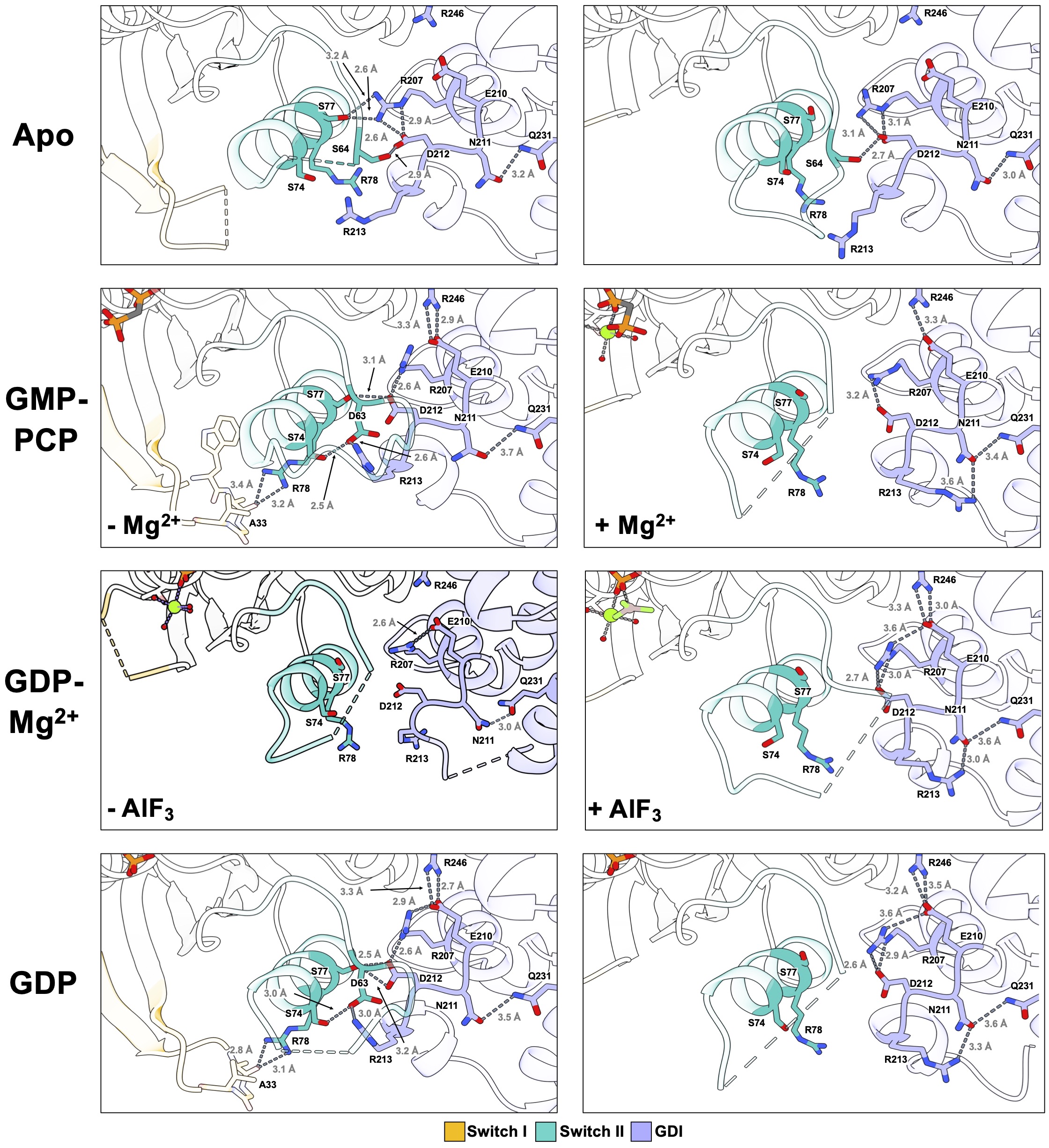

**Figure S3**. Changes in hydrogen bonding at the Switch I (yellow), Switch II (teal), and GDI (purple) sub-domains in the various states structurally characterized in this work.

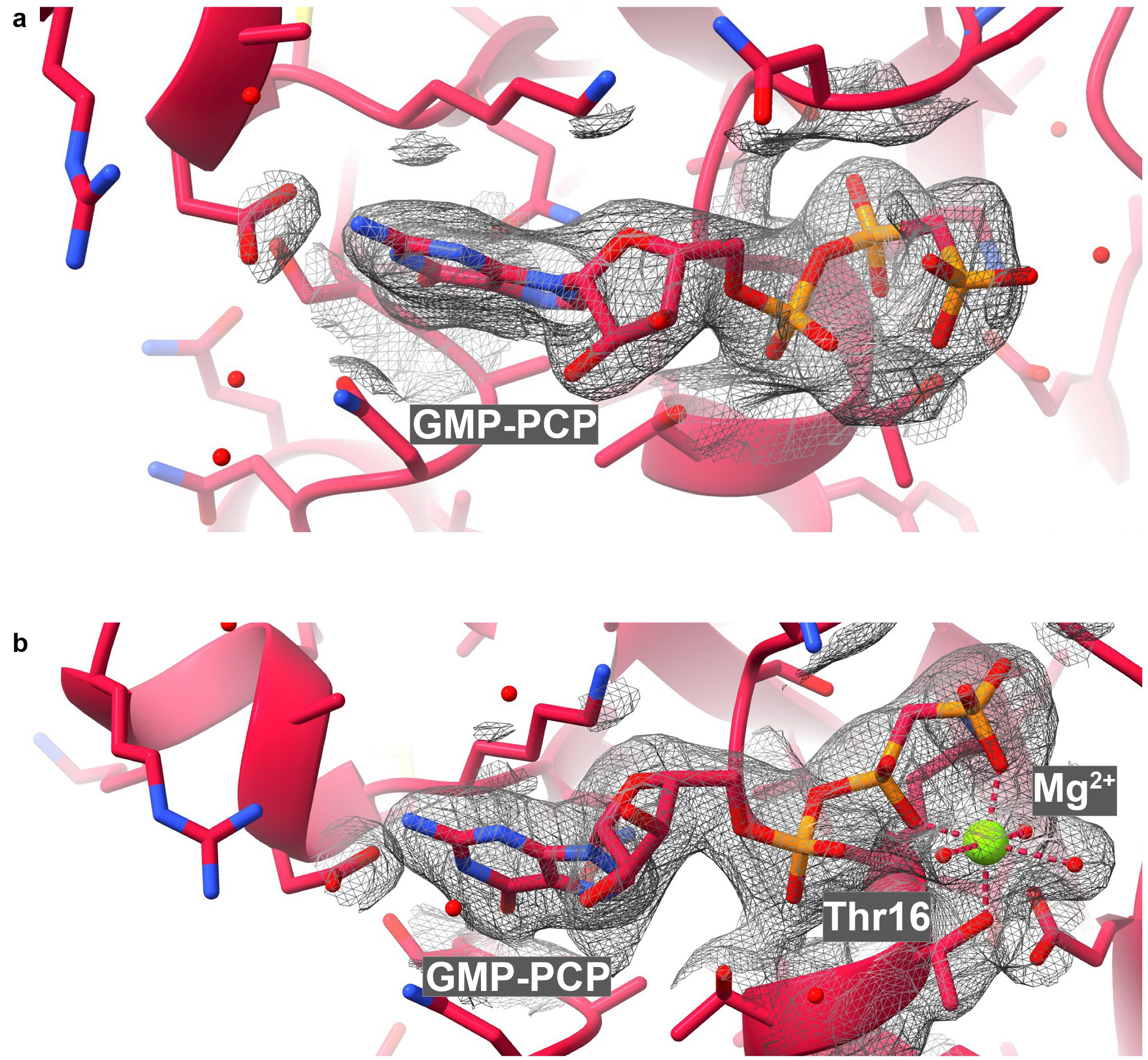

**Figure S4**. Quality of the 2*F_o_*-*F_c_* electron density surrounding the nucleotide-binding pocket contoured to 1σ (colored gray) of the GMP-PCP-bound forms of *Vc*NFeoB in the absence (**a**) and the presence (**b**) of Mg^2+^. For simplicity’s sake, only two chains of PDB ID 9D8D are shown as exemplars, chain A (**a**) and chain C (**b**).

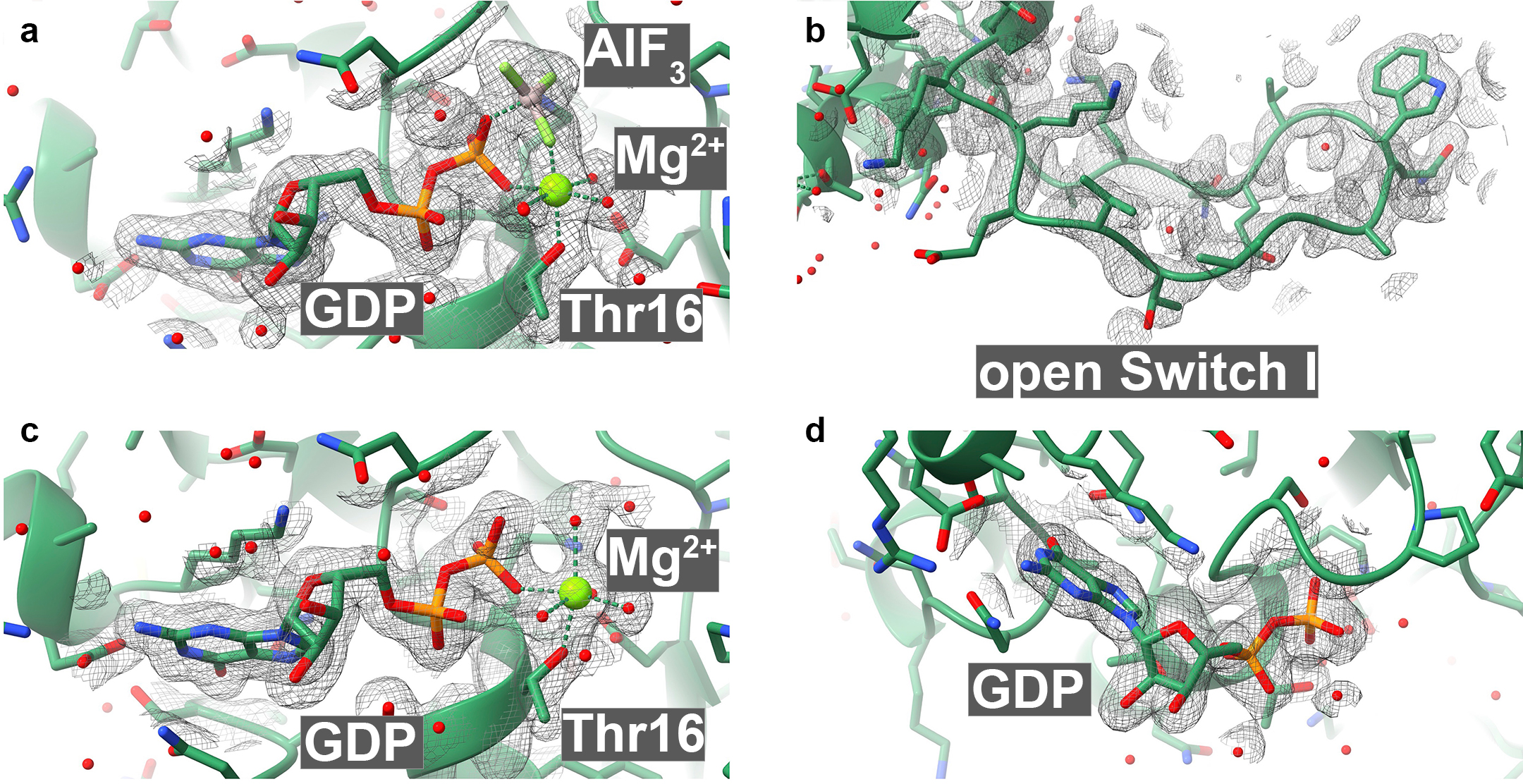

**Figure S5**. Quality of the 2*F_o_*-*F_c_* electron density surrounding the nucleotide binding pocket contoured to 1σ (colored gray) of the transition-state analog-bound *Vc*NFeoB structure. **a**. Chain B has the GDP-Mg^2+^-AlF_3_ transition-state analog bound. **b**. The Switch I region in Chain B is ordered and points away from the nucleotide-binding pocket. **c**. Chain C has the GDP-Mg^2+^ complex bound. **d**. Chains A and D have only GDP bound in the nucleotide-binding pocket. For simplicity’s sake, only Chain A of PDB ID 9PSD is shown as exemplar in this panel.

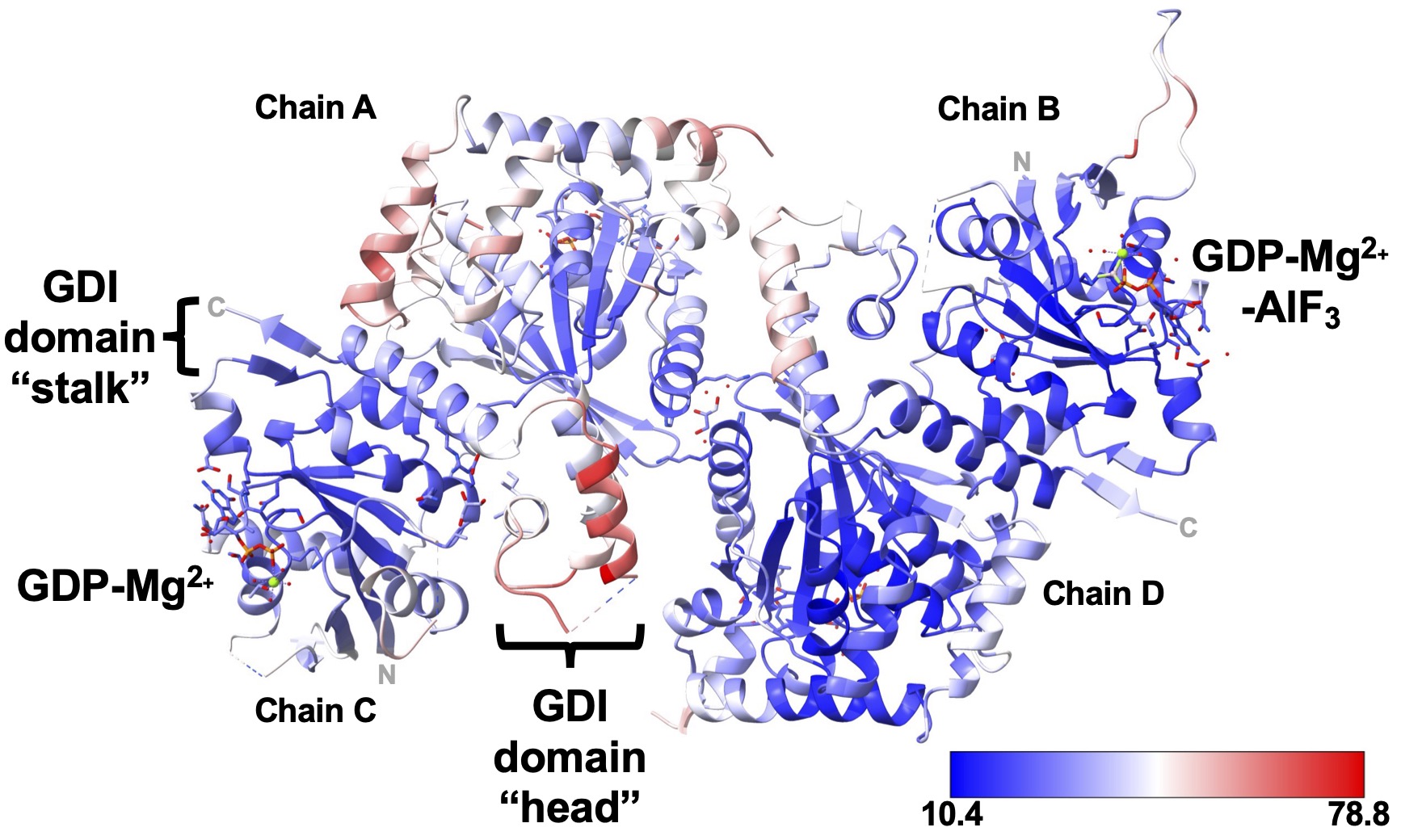

**Figure S6**. The asymmetric unit (ASU) of the transition-state analog-bound structure of *Vc*NFeoB (PDB ID 9PSD) color-coded based on B-factor value. Noteworthy are the high B-factors and local disorder within the “head” of the GDI domain in chain C (GDP-Mg^2+^-bound *Vc*NFeoB) despite its location within the center of the ASU (*i.e.*, not pointing into a solvent channel). The scale bar is color-coded from the lowest observed B-factor (10.4, blue) to the highest observed B-factor (78.8).

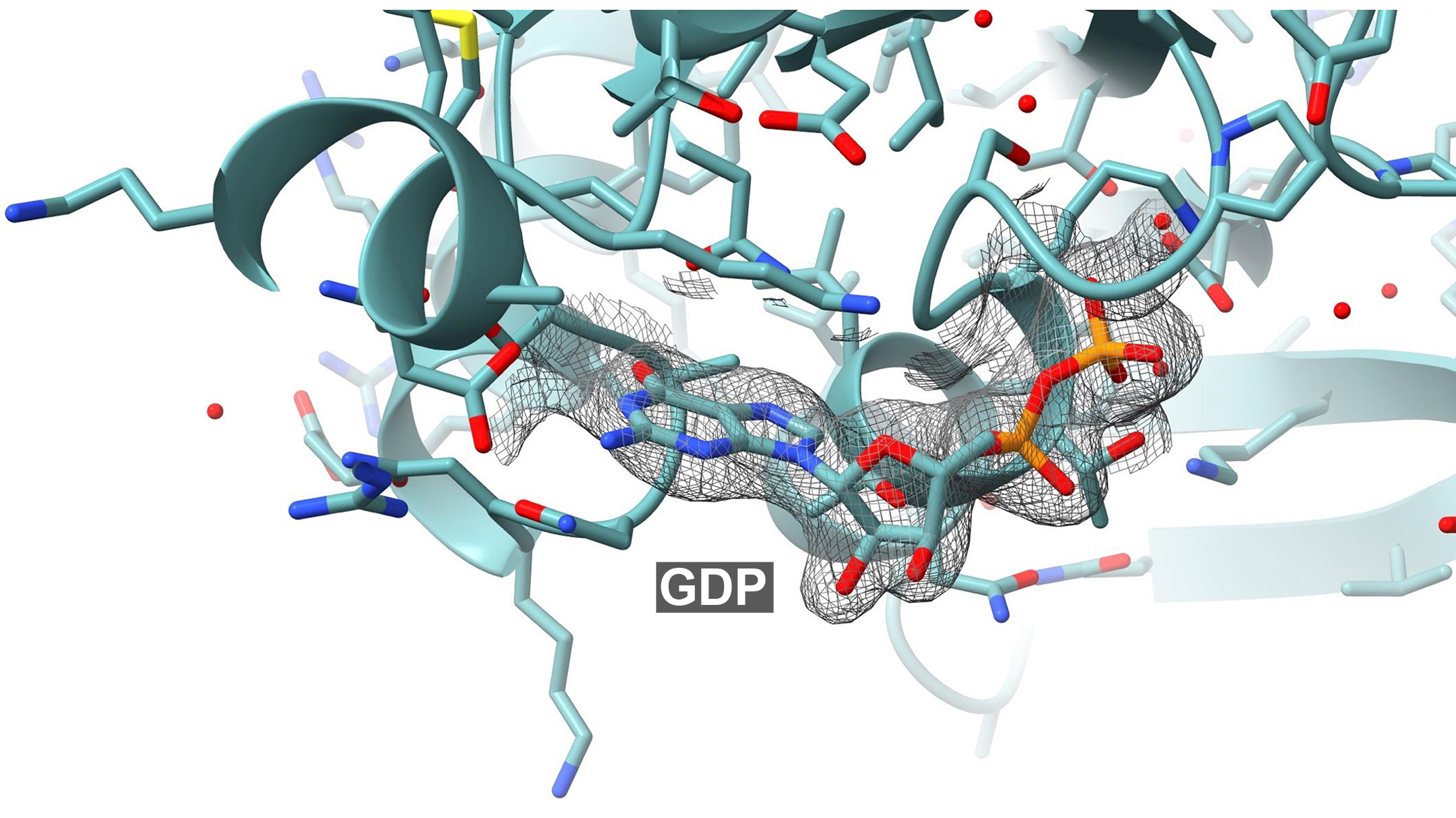

**Figure S7**. Quality of the 2*F_o_*-*F_c_* electron density surrounding the nucleotide binding pocket contoured to 1σ (colored gray) of GDP-bound *Vc*NFeoB structure. For simplicity’s sake, only Chain A of PDB ID 9D8B is shown here as an exemplar.
